# The meninges host a unique compartment of regulatory T cells that bulwarks adult hippocampal neurogenesis

**DOI:** 10.1101/2024.06.17.599387

**Authors:** Miguel Marin-Rodero, Elisa Cintado Reyes, Alec J. Walker, Teshika Jayewickreme, Felipe A. Pinho-Ribeiro, Quentin Richardson, Ruaidhrí Jackson, Isaac M. Chiu, Christophe Benoist, Beth Stevens, José Luís Trejo, Diane Mathis

## Abstract

Our knowledge about the meningeal immune system has recently burgeoned, particularly our understanding of how innate and adaptive effector cells are mobilized to meet brain challenges. However, information on how meningeal immunocytes guard brain homeostasis in healthy individuals remains sparse. This study highlights the heterogeneous and polyfunctional regulatory-T (Treg) cell compartment in the meninges. A Treg subtype specialized in controlling Th1-cell responses and another known to control responses in B-cell follicles were substantial components of this compartment, foretelling that punctual Treg-cell ablation rapidly unleashed interferon-gamma production by meningeal lymphocytes, unlocked their access to the brain parenchyma, and altered meningeal B-cell profiles. Distally, the hippocampus assumed a reactive state, with morphological and transcriptional changes in multiple glial-cell types; within the dentate gyrus, neural stem cells showed exacerbated death and desisted from further differentiation, associated with inhibition of spatial-reference memory. Thus, meningeal Treg cells are a multifaceted bulwark to brain homeostasis at steady-state.

**One sentence summary:** A distinct population of regulatory T cells in the murine meninges safeguards homeostasis by keeping local interferon-γ-producing lymphocytes in check, thereby preventing their invasion of the parenchyma, activation of hippocampal glial cells, death of neural stem cells, and memory decay.

---

The meninges constitute a three-layered structure just under the skull and vertebral column, covering the brain and spinal cord. This brain border, in particular the dura mater layer, hosts a dense and highly diverse constellation of immunocytes at homeostasis (*1, 2*) as well as a dedicated lymphatic drainage system (*3*). Many of these cells are members of the innate immune system, above all macrophages (MFs). But there are also small populations of lymphocytes whose functions are only beginning to be understood. For example, severe combined immunodeficiency patients, as well as mice lacking T and B cells, have behavioral abnormalities that resolve with reconstitution of the adaptive immune system (*4–7*). T cells [α:β T, γ:δ T and mucosal associated invariant T (MAIT) cells] seem especially important for brain homeostasis because the cytokines they produce [e.g. interleukin (IL)-4, IL-10, interferon (IFN)γ, tumor necrosis factor (TNF)α, and IL-17] impact various behavioral parameters as well as promote cognitive changes with aging (*8–17*).

Even sparser is information on cellular regulators of meningeal immunocyte responses. Most types of immune reactions are controlled by Foxp3+CD4+ regulatory T (Treg) cells (*18*). These cells can control an immune response directly by acting as a sink for IL-2, by secreting suppressive factors like IL-10 or transforming growth factor (TGF)ß, or by expressing co-inhibitory molecules such as CTLA-4, PD-1, or LAG-3. In addition, they can act indirectly by modulating the differentiation and presentation capacity of antigen-presenting cells (APCs) such as dendritic cells (DCs) and MFs. Treg cells in non-lymphoid tissues (termed “tissue-Tregs”) also control non-immunological processes, notably promoting tissue repair in response to injury, including an impact on central nervous system (CNS) pathologies such as stroke or experimental autoimmune encephalomyelitis (EAE) (*19*).

Treg-cell activities have been studied in the meninges of old or diseased mice (e.g. (*20*)), but here we have addressed their role in promoting brain health, i.e. in immunological homeostasis of the brain meninges and parenchyma. Our work highlights a little-known population of Tregs in the meningeal dura mater of healthy mice, clustered with DCs along the sinuses. These Treg cells rein in local inflammation, in particular IFNγ production, and shield the brain’s functional integrity at steady state, preventing immunocyte infiltration into the parenchyma and preserving neurogenesis within the hippocampal niche. Our findings both extend the purview of tissue-Treg cells and bring to light an important immunological regulator of brain homeostasis.

## The dural layer of the meninges hosts a population of Foxp3+CD4+ T cells at steady-state

While our knowledge of the immunocyte populations operating in the meninges – particularly mouse meninges – has greatly improved over the past several years (*2*), we still know very little about the representation, phenotype, and function of meningeal Treg cells, especially at steady-state. To begin filling this knowledge-gap, we performed a cytofluorometric analysis of the Foxp3+CD4+ compartment of meninges isolated from transcardially perfused, 6-week (wk)-old, male, C57BL/6 (B6) mice (see Fig. S1A for the gating strategy). We chose to focus on the dura mater because the other meningeal layers contain far fewer immunocytes at steady state (*2, 21*), and because these layers are substantially more difficult to access, thereby greatly prolonging isolation time and thus increasing the risk of stress and cell damage. As illustrated in Fig. 1A, we could readily identify a population of Foxp3+CD4+ T cells in the meninges, fractionally slightly higher than in the spleen (Fig. S1B), greater than an order of magnitude more abundant then in the leptomeninges (>15x), and much higher than in the choroid plexus (>29x) (Fig.S1C). At an average of about 125 Treg cells per mouse, the meningeal Treg population was evident but small enough to limit some of the experimental approaches we could eventually pursue.

**Figure 1.**
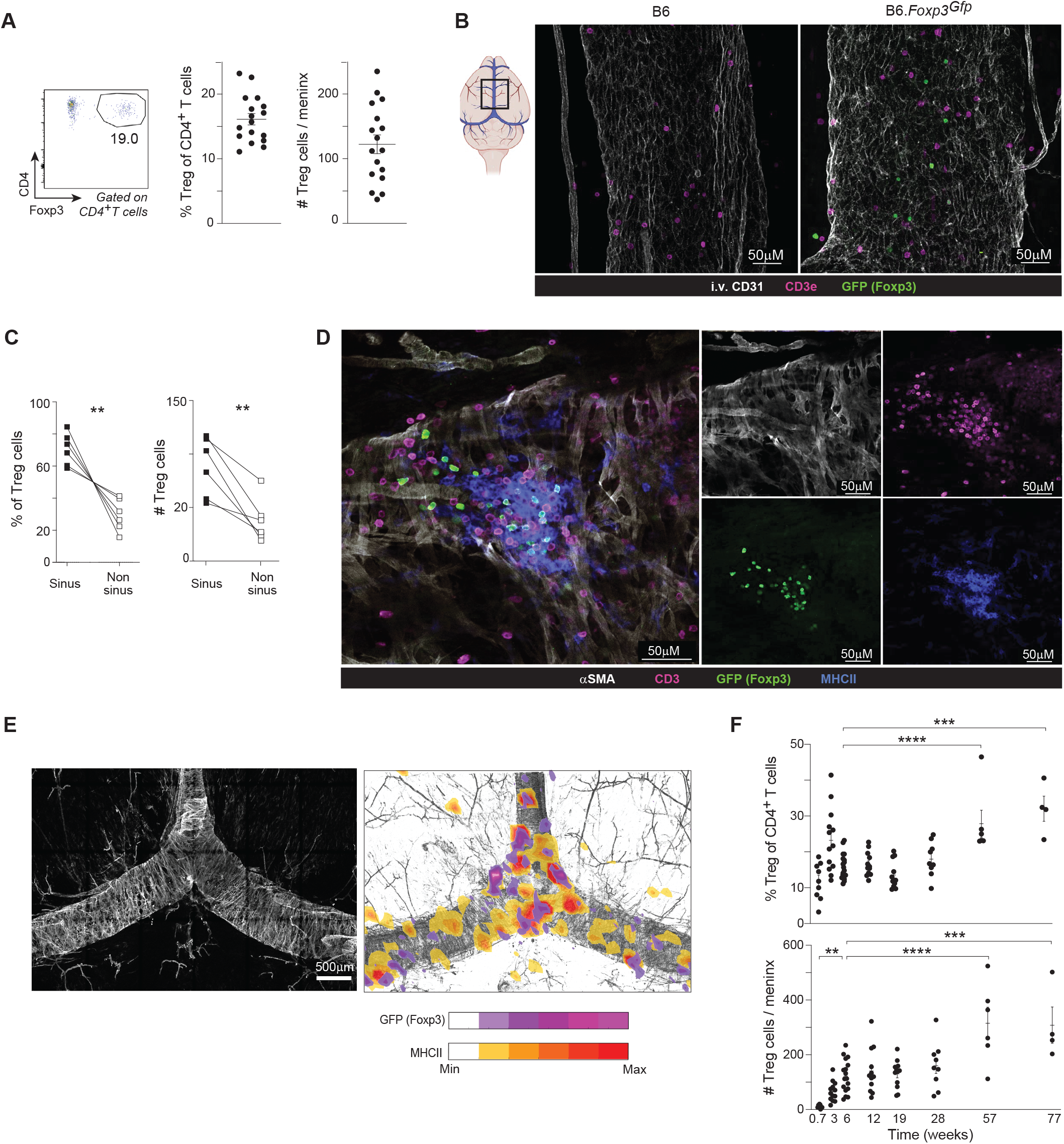
A population of Foxp3+CD4+ T cells within the meningeal dura at steady-state. (A) Flow cytometry of Foxp3+CD4+ T cells in the dural meninges of 6wk-old mice. Left panel: representative flow-cytometric plot; Right panels: summary data. n=17 (B) Confocal imaging of transverse sections of the dura mater. Left panel: graphical representation of the meningeal sinuses, with the region of interest delineated by a square; right panels: representative images. (C) Representative confocal image of a meningeal Treg-cell cluster on a transverse section stained for the indicated marker proteins. (D) Quantification of Treg-cell locations. n=6 (E) Tile-scan images of multiple duras registered to a reference map (left), plotting average densities of the Foxp3-GFP and MHC-II signals across 6 tissues (right). (F) Quantification of Treg cells in the dura mater across time. n ≥4 Each data-point is from an individual mouse. iv, intravenous; SMA, smooth-muscle actin; MHC, major histocompatibility complex. Mean ± SEM. ∗p < 0.05, ∗∗p < 0.01, ∗∗∗p < 0.001.

Confocal microscopy of tissue obtained from B6.*Foxp3Gfp* mice provided an independent confirmation of the meningeal Treg compartment. Most Treg cells were close to the dural sinuses, accompanied by other types of T cells (CD3e+) (Fig. 1B). While some Treg cells were dispersed along the sinuses, they often formed clusters, especially with other T and major histocompatibility complex (MHC)II+ cells (Fig. 1E, F). The latter cells were mostly type-2 conventional DCs (cDC2s), defined by the marker profile MHCII+CD11c+F4/80-Sirpa+ (*22*) (Fig. S1D, E). The clustering of Treg and MHCII+ cells within the dural sinuses was confirmed in multiple mice ranging from 5 to 16 wks of age (Fig. S2A, B), best visualized in Fig. 1E, where, in order to gain a more comprehensive view of the localization of Treg cells and APCs in the dura mater, we registered tile-scan images of multiple duras to a reference map and then plotted the average density of GFP (Foxp3) and MHCII signals across all 6 tissues.

Aging was associated with a decline in the total number of CD45+ cells in the meninges (Fig. S2C); in contrast, the fraction and number of meningeal Treg cells showed a general increase with age (Fig. 1F), consistent with a published comparison of 2- and 24-month-old mice (*20*). Total CD45+ cells and Treg cells accumulated in the meninges of male and female mice to a similar degree (Fig. S2D, E).

## The meningeal Treg compartment is a heterogeneous population with repeated T cell receptor (TCR) sequences

Treg cells operating in non-lymphoid tissues have distinct transcriptomes, TCR repertoires, and growth-factor dependencies, adapted for optimum function and survival within their particular settings (*19*). As a first step in exploring the phenotype and functional potentials of meningeal Treg cells, we performed population-level RNA sequencing (RNA-seq) on three replicates, each pooled from five 15wk-old males. Principal Component Analysis revealed the Treg cells in meninges to be distinct from those of lymphoid tissues and all other non-lymphoid tissues, including inflamed brain (*23*) (Fig. S3A). Nonetheless, meningeal Tregs preferentially expressed a previously reported pan-tissue-Treg signature (*19*) in comparison with their splenic counterparts (Fig. S3B). Pathway analysis revealed the meningeal-Treg transcriptome to be enriched in transcripts encoding molecules involved in cell adhesion and locomotion (e.g. *Ccl5*, *Cxcl10*, *Itga4*, *Itgav*, *Ccr2*, *Cxcr3*); promotion of cell death; the inflammatory response; regulation of myeloid-cell differentiation; and, potentially most relevant here, regulation of neuron death (*Ifng*, *Il10*, *Cd200r*, *Wisp*, *Ptgs1*) (*24–30*) (Fig. S3C). The transcriptomes of meningeal Treg cells from male and female mice were very similar, the major difference being an enrichment for cell-cycle pathways in males (Fig. S3D, E), a finding confirmed by their higher fraction of Ki67+ cells (Fig. S3F). Similarly, an elevation in cell-cycle pathways was the major transcriptional feature distinguishing adult (16wk-old) from old (53wk-old) mice (Fig. S3G, H).

Skull bone-marrow (BM) and the meninges are interconnected by small conduits through which myeloid and B cells are able to migrate (*31–33*). Although T cells have been reported to access the meninges via the circulation (*32, 34*), we compared the transcriptomes of Treg cells isolated from the meninges and skull BM in part to assess potential contamination of the meningeal-Treg isolate by skull-BM cells. The two Treg compartments were clearly different (Fig. S3J), the meningeal population being relatively impoverished in transcripts specifying molecules related to catabolic processes and protein transport and enriched in transcripts encoding proteins involved in metabolic pathways (Fig. S3I).

It is by now well established that multiple Treg subtypes exist, specialized to perform diverse functions at diverse sites (*19*), and that tissue-Treg compartments are actually a conglomeration of multiple subtypes (*35–37*). To explore the heterogeneity of meningeal Treg cells, we flow-cytometrically sorted the total CD4+ T cell populations from two cohorts of mice (15 and 25 individuals) and performed scRNA-seq coupled with sc*Tra/b*-seq. After quality control, we retained a total of 4090 CD4+ T cells, with a mean of 2453 unique fragments per cell. Dimensionality reduction, unsupervised clustering, and signature overlay revealed a clear group of *Foxp3+* Treg cells (Fig. S4A), which we extracted for downstream analyses. Re-clustering the Treg cells revealed three distinguishable subtypes (Fig. 2A), each with approximately equal fractional contributions from the two scRNA-seq replicates (Fig. S4B). A heatmap of the most differentially expressed genes confirmed the cluster parsing (Fig. 2B), and the genes’ identities prompted us to designate the clusters “T helper (Th)-1-like”, “T follicular regulatory (Tfr)- like”, and “quiescent.” Th1-like Treg cells express the transcription factor (TF) Tbet and multiple interferon-stimulated genes (ISGs), notably *Cxcr3*, and are adept at controlling Th1, CD8+ T, and NK cell responses (*38, 39*). Tfr cells, identified by expression of transcripts encoding CXCR5 and PD-1, are generally localized in germinal centers (GCs), where they control the magnitude and output of the local B-cell response, dampening production of autoantibodies (*40, 41*). The quiescent cluster, typical of tissue-Treg cells, expresses a number of genes characteristic of unreactive Treg cells (*36, 37*). The three subtype designations were further solidified by overlaying corresponding gene signatures (*19, 36, 42, 43*) (Fig. 2C) or key marker transcripts (Fig. S4 C-E) on the UMAP. Lastly, flow cytometry confirmed the existence of the three subtypes within the meningeal Treg compartment, each as a higher fraction of CD4+ T cells than was found in in the spleen (Fig. 2D-F). Also consistent with the Treg subtype designations is that analogous processing of the non-Treg CD4+ T cells from the same scRNA-seq cohorts yielded clear Th1, IFN-responsive, and Tfr clusters (Fig. S4F).

**Figure 2.**
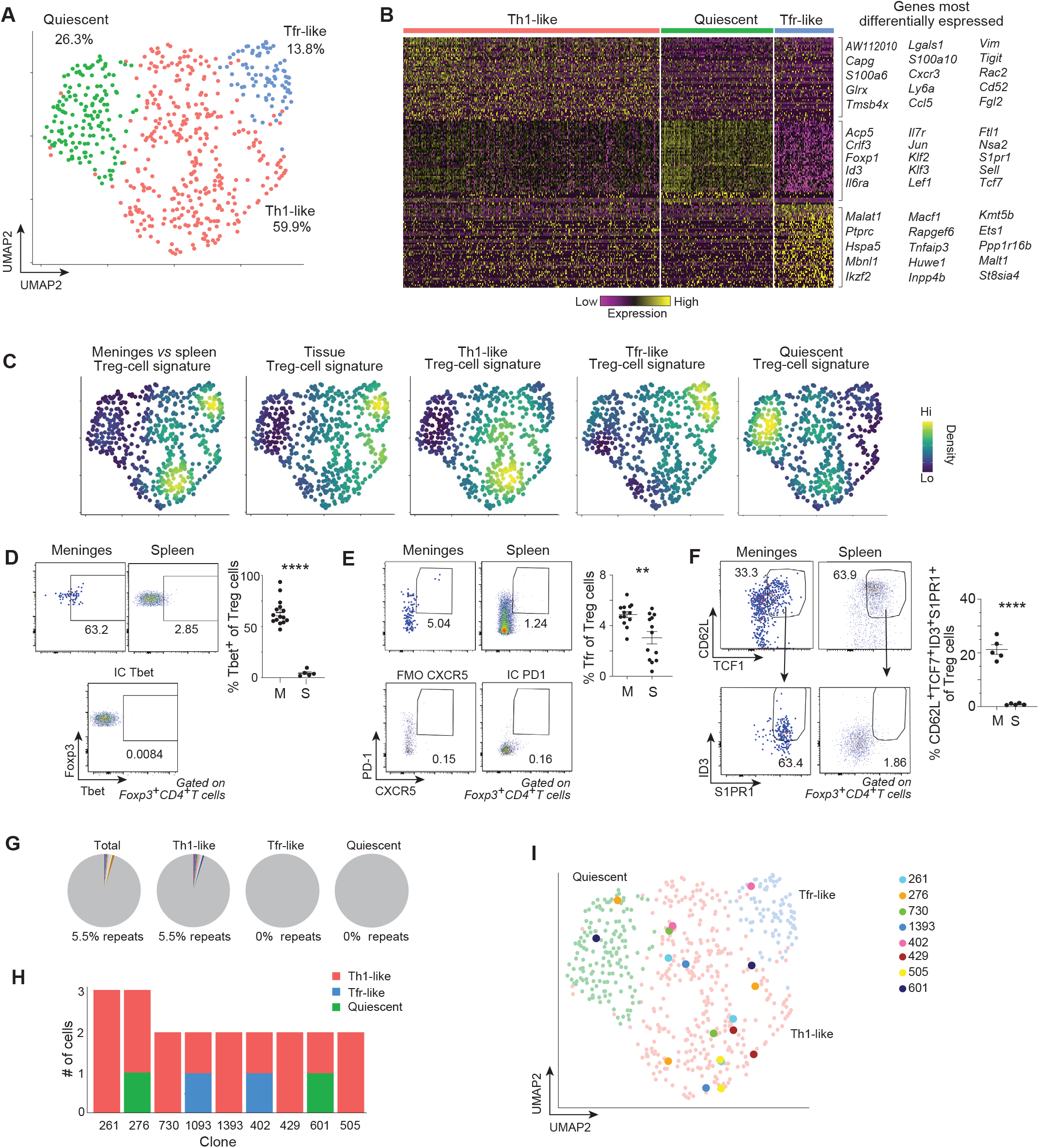
Heterogeneity of and clonal expansions within the meningeal Treg compartment. (A-C) Merged scRNA-seq data from two cohorts of dura mater Treg cells, 15 and 36wks old (n=40). (A) UMAP plot. Percentages indicate the fractional representation of each cluster. (B) Heatmap of the 50 genes most differentially expressed by each cluster. The 10 genes (excluding *Rps* genes in cluster 2) most specific are indicated to the right. (C) Expression density plots of the indicated signatures. Signature references are in the Methods section. (D-F) Flow cytometry of the three meningeal Treg-subtypes’ key marker proteins. Left: representative dot plots; right: summary data. On E and F the representative flow plots depict 3 samples concatenated. (G-I) sc*Tcr*-seq data from the meningeal Treg-subtypes. (G) Pie-charts showing the proportion of clonally expanded cells in each cluster. Individual clones are depicted by different colors; non-expanded clones are in gray. (H) Bar plot showing the clonal overlap between the various clusters. x-axis: clone names: y-axis: number of Treg-cells of that particular clone. Bar colors correspond to the clusters of panel A. (I) Expanded TCR clones situated on the UMAP plot from panel A. UMAP, Uniform Manifold Approximation and Projection; Tfr, T follicular regulatory cell; Th1, T helper1; M, meninges; S, spleen. Mean ± SEM, p-values as per Fig. 1.

From the same two pools of meningeal Treg cells, we also obtained sc*Tra/b* sequences. We were especially interested in “repeat-sequences,” defined as the presence of two or more cells with the exact same nucleotide sequences encoding both their TCRα and TCRβ chains, suggestive of clonality. Even though we could examine sequences from a total of only 366 cells from a total of 40 mice (Table S1, GSE234317), making detection of repeats rather unlikely, we observed 5.5% repeats within the total dataset, intra-subtype sharing being restricted to the Th1-like cluster (Fig. 2G). In addition, a few repeats were shared between the Th1-like cluster and the Tfr-like or quiescent clusters (Fig. 2H, I).

## Meningeal Treg cells rein in local lymphocytes, in particular their production of IFNγ

Punctual Treg-cell depletion was accomplished in 6wk-old B6.*Foxp3.Dtr+* and B6.*Foxp3.Dtr--* male littermates (hereafter referred to as DTR+ and DTR-, respectively) by intraperitoneal (ip) injection of diphtheria toxin (DT) on three consecutive days, followed by analysis 3 days after the last injection (Fig. 3A). This relatively short depletion protocol, chosen to minimize long-range systemic influences, effectively reduced the representation of Treg cells in the meninges (Fig. 3B) without significantly augmenting total immunocyte numbers (Fig. 3C). Nonetheless, the fractions and numbers of αβT and B cells increased and decreased, respectively, with no evident changes in the representations of other major immunocyte types (Fig. 3D). Within the T cell population, both the CD4+ and CD8+ fractions expanded at the expense of the double-negative fraction (Fig. 3E). Confocal imaging of meninges whole-mounts revealed that the expanded T-cell populations were confined to the typical niche, accompanied by an increased MHCII signal (Fig. 3F). The B-cell loss reflected a reduction in the follicular B cell stages (Fig. 3G; Fig. S5A for the gating strategy). Meningeal Treg cells formed clusters in close proximity to B cells (Fig. 3H).

**Figure 3.**
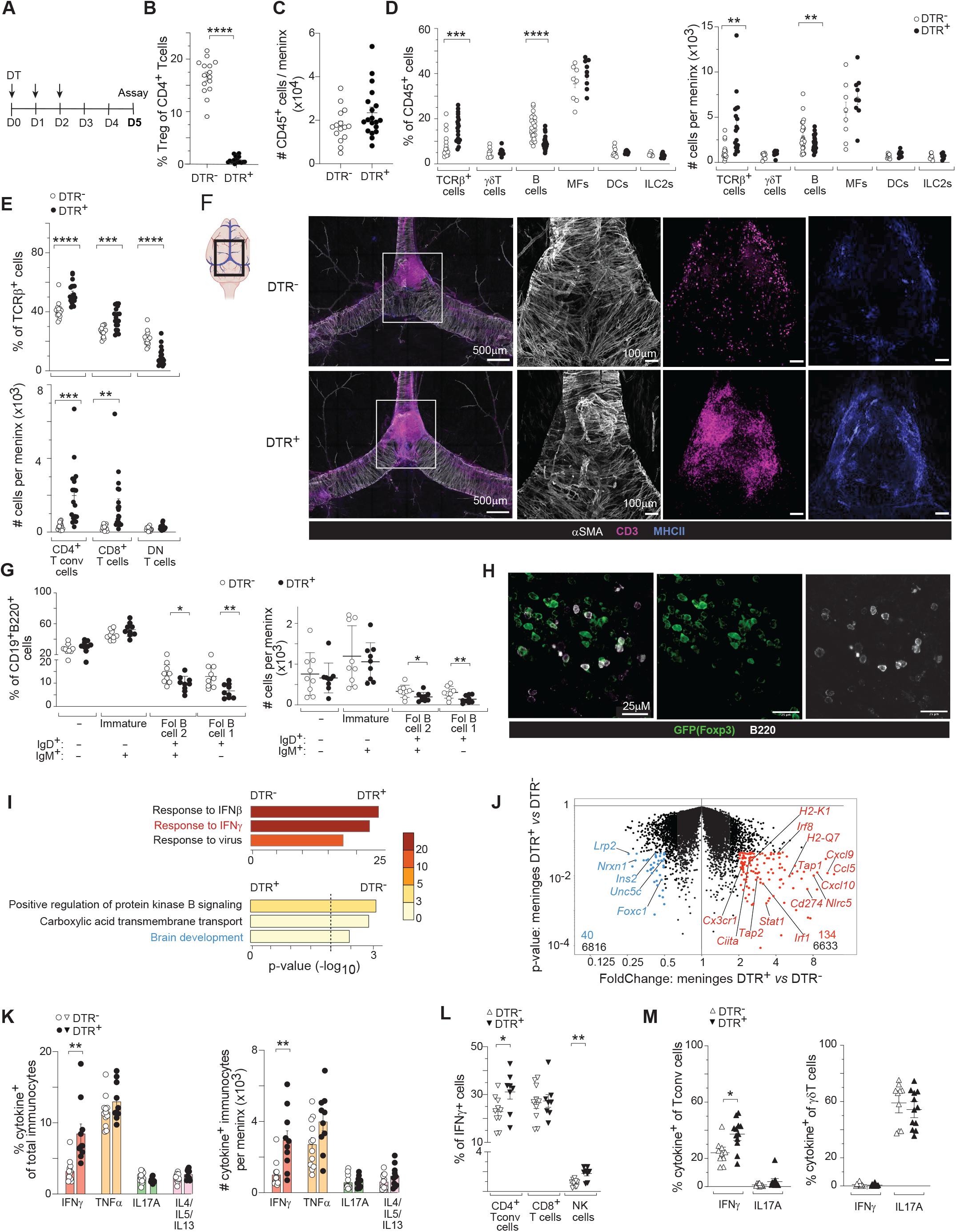
Influences of Treg cells on meningeal lymphocyte. (A) Treg-cell depletion protocol. 6wk-old male DTR*+* and DTR*-* littermates were ip-injected with DT as indicated, and assays were performed on day 5 (D5) after the first injection. (B) Percentage of meningeal Treg cells. n ≥15 (C) Total meningeal immunocyte counts. n ≥15 (D) Percentage (left) and numbers (right) of select immunocyte populations. n≥8 (E) Percentage (upper) and numbers (bottom) of the major T cell subsets. n≥15 (F) Imaging of meningeal sinuses from DT-treated DTR- and DTR+ mice. Graphical representation showing the region of interest is shown to the left. Then a representative confocal image of the three indicated strains. Then higher power images of the white-squared region stained with, in order: αSMA for sinus structure, αCD3 for T cells and anti-MHCII for antigen presenting cells. (G) Percentage (left) and numbers (right) of meningeal B cell populations. n=9 (H) Representative immunofluorescence image of Treg and B cells in the meninges. (I) Pathway enrichment analysis via Metascape (*131*) of the genes differentially expressed in panel J [fold-change (FC) >1.5, p-value <0.05]. (J) Volcano plot of population-level RNA-seq comparing meninges in the presence and absence of Treg cells. Annotated transcripts are those involved in the pathways highlighted in Fig. 3I (interferon signaling and brain development pathways). Triplicate samples. (K) Percentage (left) and numbers (center) of cytokine positive immunocytes from the dura mater. (L) Representation of the IFNγ producers from (K). (M) Percentage of major cytokine producers from the dura mater. DT, diphtheria toxin; DTR, DT receptor; MF, macrophages; NF, neutrophils; DCs, dendritic cells; ILC, innate lymphoid cells; Fol, follicular; other abbreviations as per Fig. 1. Mean ± SEM. ****, p<0.0001; other p-values as per Fig. 1.

Since immunocytes represent only around 40% of live cells in the meninges, we examined the Treg-depleted meningeal environment more comprehensively by performing whole-tissue RNA-seq of DTR- and DTR+ littermates 3 days after the last DT injection. Meninges lacking Treg cells were enriched in IFN-response signatures (both type I and II), while being slightly impoverished in pathways related to brain development (e.g. lower expression of *Lrp2, Nrxn1, Ins2, Unc5c, Foxc1*) (Fig. 3I, J). For confirmation of IFN induction, and to identify culprit IFN-producing cells, we cytofluorometrically quantified meningeal cells expressing IFNγ and other lineage-defining cytokines from DTR- and DTR+ mice (see Fig. S5B for the gating strategy). After Treg-cell depletion, immunocytes expressing IFNγ, but not those making the other cytokines examined, were significantly enriched in the meninges (Fig. 3K). Foxp3-CD4+ conventional T (Tconv) cells and CD8+ T cells were the major producers of IFNγ in both the presence and absence of Treg cells; loss of Treg cells induced further IFNγ production by CD4+ Tconv and NK cells, although the contribution of NK cells remained low (Fig. 3L). Neither the fraction of Th17 cells nor that of IL-17-producing γδT cells was increased in the meninges in the absence of Treg cells (Fig. 3M).

To address whether it was Treg cells specifically in the meninges that drove these effects, we performed two types of experiments. First, we injected an anti-CD25 monoclonal antibody (mAb) or an irrelevant anti-IgG control mAb into the intracisternal magna and performed whole-tissue RNA-seq 72 hours later. Anti-CD25 injection has been a frequently used alternative method to deplete Treg cells (at least those that are CD25+) (e.g. (*44–46*)); and intracisternal injection allows preferential delivery to the meninges while minimizing systemic spread (but technical difficulties permit only a single injection). This protocol, as previously described, did not change total Treg cell numbers but completely eliminated the CD25+ Treg component (determined using a different, non-competing anti-CD25 mAb) (Fig. S5C). Nonetheless, this method of Treg-cell ablation also elicited a clear response to IFNγ (Fig. S5D, E). Second, exploiting up-regulation of MHCI as a well-accepted marker of IFNγ sensing, we compared the time-course of IFNγ responsiveness in the meninges and spleen (Fig. S5F, G). In the meninges, up-regulation of MHCI expression was evident on both immunocytes and stromal cells by 24 hours after the first DT injection, while in the spleen, up-regulation was observed only at 72 hours [consistent with (*47*)] and only on stromal cells (Fig. S5H). Along with the short depletion protocol we employed, these two sets of findings argue that the punctual ablation of meningeal Treg cells incited a local response, and the resulting immunological changes did not simply reflect systemic effects issuing from loss of the circulating Treg pool.

## Treg cells guard against lymphocyte invasion of the brain parenchyma

Over the past several years, it has become clear that immunocytes within the dura mater, especially T cells, have a direct route of communication with the brain parenchyma (*2*). Moreover, it is now recognized that the meninges are indispensable for proper brain function (*48*), with intrinsic roles in B cell differentiation (*49, 50*), innervation (*51*), and defense against pathogens (*51, 52*). Thus, an obvious question was what happens in the brain upon punctual Treg-cell depletion? So we depleted Treg cells as per Fig. 3A, and performed flow-cytometric, immunohistologic, and transcriptomic comparisons of the brain parenchyma from wild-type (DTR-) and Treg-deficient (DTR+) littermates.

More CD45+ cells – particularly αβT and NK cells – were found in the brain parenchyma of perfused DTR+ than DTR- mice (Fig. 4A, B). There seemed to be no preference for CD8+ vs CD4+ T cells (Fig. S6A, B). Residual contamination by circulating cells was identified by intravascular labelling of blood cells with a fluorescent anti-CD45 mAb (*53*), revealing that just 9.7% and 12.1% of CD45+ cells in the parenchyma and meninges, respectively, were circulating at steady-state (Fig. S6C). After Treg-cell depletion, the brain parenchyma from DTR+ mice had more circulating T, but not NK, cells than did parenchyma from their DTR- littermates, opposite to what was observed in the meninges (Fig. S6D-E).

**Figure 4.**
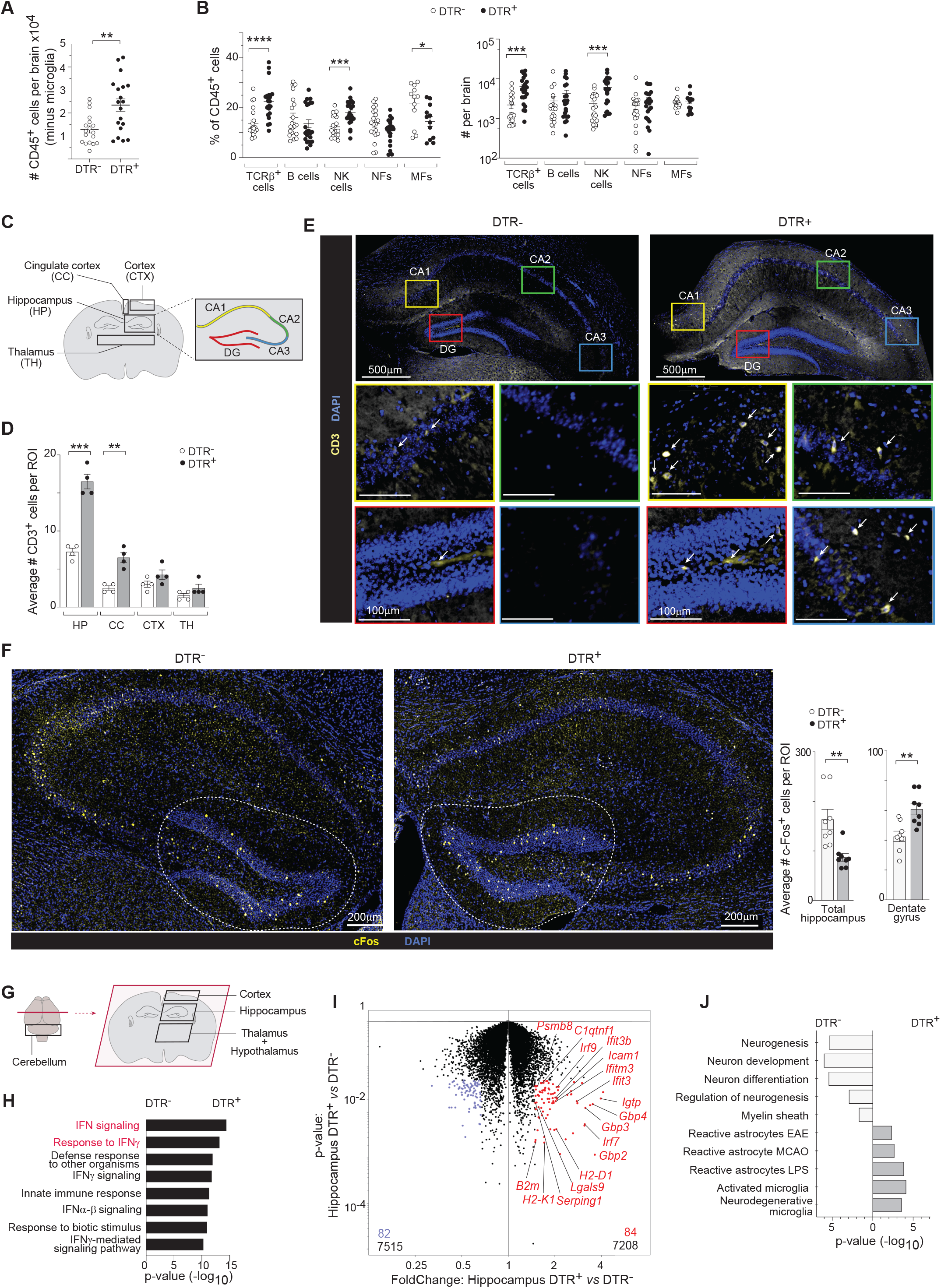
T cell invasion of the brain parenchyma in the absence of Treg cells. (A-B) Flow-cytometric analysis of the brain parenchyma of mice depleted of Tregs or not, as per Fig. 3A. Males aged 6 to 7 wks old. (A) Total immunocyte numbers. n ≥ 18 (B) Frequency (left) and numbers (right) of select immunocyte populations. n ≥ 11 (C-E) T cell invasion into the brain parenchyma after depletion of Treg cells in DT-treated DTR+ mice, as per Fig. 3A. (C) Diagrammatic representation of the brain with regions of interest delineated. Coronal section. (D) Quantification of the average CD3+ T cells as described in the Methods section. n= 4 (E) Representative confocal images of coronal sections of the whole hippocampus. Top rows: regions of interest. The quartet of higher power images below show anti-CD3 mAb (+DAPI) staining for T cells in each region of interest, framed in the colors corresponding to those of the top panel. Arrows point to T cells. (F) Left: representative confocal images of the hippocampus’ cFos staining pattern in the presence (left) or absence (right) of Treg cells. The dotted line delimits the DG. Right: quantification of cFos+ cells in the hippocampus and DG as described in the Methods section. n≥8 (G-J) Whole-tissue RNA-seq analysis of brain regions from mice depleted of Treg cells or not, as per Fig. 3A. (G) Diagram showing regions of interest. (H) Pathway-enrichment analysis via Metascape (*131*) of the differentially expressed transcripts from panel I [FC>1.5; p-value<0.05]. (I) Volcano comparing DTR+ and DTR- hippocampus. Annotated genes are those involved in pathways highlighted in H (interferon signaling and response to IFNγ). Triplicate samples. (J) Specific analysis via Gene Ontology (GO) and GSEA (MsigDB) database of pathways related to glial activation and hippocampal functions. DG, dentate gyrus; CA, Cornu Ammonis; other abbreviations as per Fig. 3. Mean ± SEM, p-values as per Fig. 3.

To localize the T cells infiltrating the brain, we first performed confocal microscopy of anti-CD3-stained sections from four brain regions: the hippocampus, cingulate cortex, cortex region overlying the hippocampus, and thalamus (Fig. 4C). Ablation of Treg cells resulted in significantly more T cell accumulation in the hippocampus and, to a lesser extent, in the cingulate cortex but not in the other two brain regions examined (Fig. 4D). Infiltrating T cells were scattered throughout the hippocampus (Fig. 4E). A second approach was to stain brain sections with a mAb recognizing cFos, an immediate-early marker of cell activation and thereby neuronal activity (*54*). cFos+ cells were reduced in the hippocampus as a whole and in the cortex just above it but were specifically increased in the dentate gyrus (DG) of the hippocampus, the site of adult neurogenesis (Fig. 4F; Fig S6F). For a comprehensive view of brain changes subsequent to Treg-cell depletion, we performed whole-tissue RNA-seq on four brain regions: the cortex, hippocampus, thalamus plus hypothalamus (T&H), and cerebellum (Fig. 4G). Primarily in the hippocampus (Fig. 4H, I), but to a lesser extent in the T&H and cortex regions as well (Fig. S6G, H), there was an enrichment for IFNγ-signaling pathways. In stark contrast, the cerebellum did not show an enrichment in these gene-sets (Fig. S6G, H), even though transcripts encoding IFN receptors were amply expressed throughout the brain (Fig. S6I), arguing that induction of the IFNγ-signaling pathways was not simply a generalized systemic effect. Consistent with this important point, a time-course comparison of the transcriptional response to punctual loss of Treg cells in the meninges vs hippocampus revealed the former to significantly precede the latter, as evidenced by the splaying of both total and IFNγ-induced transcripts along the x-axis of the 12hr FoldChange (FC)/FC plots of Fig. S6J.

We also specifically addressed whether one of the main functions of the hippocampus, adult neurogenesis (*55*), was altered in the absence of Treg cells. Indeed, the hippocampus of DTR+ mouse brains showed an enrichment in gene signatures indicative of reactive and disease-associated glia (*54, 56–59*), coupled with an impoverishment in signatures associated with myelination, neuronal differentiation, and neurogenesis (*Gfap, Sox2, Ascl1, Hes1, Wnt2)* (Fig. 4J, Fig. S6K).

## Treg cells bridle glial cells in the hippocampus

Glial cells, both microglia and astrocytes, are vital for brain formation and function (*60*). In the hippocampus, specifically, microglia are important for the removal of newborn neurons undergoing apoptosis and for regulation of the survival and proliferation of new neurons (*61, 62*). A brain insult provokes microglia to react rapidly, changing their morphology from a ramified to an ameboid form (*63*), but their over-reaction can be detrimental for neuronal survival and neurogenesis (*64, 65*). Hippocampal astrocytes are key regulators of neuronal stem-cell proliferation and differentiation (*66*) and of memory formation (*67–69*).

To explore the state of glia after punctual Treg-cell depletion (as per Fig. 3A), we first performed confocal microscopy on hippocampal (dentate gyrus) sections stained with cell-type-specific mAbs. Iba1+ microglia acquired the morphology of activated cells, with reduced arborization, evidenced by fewer branches and processes per cell, fewer branches per process, and shorter processes (Fig. 5A). The high abundance and low heterogeneity of adult hippocampal microglia (*70*) permitted us to sort them from DTR- and DTR+ littermates and perform population-level RNA-seq. Superimposing disease-associated and damage-associated microglia signatures on volcano plots of transcripts from microglia sorted from mice of the two genotypes revealed significant over-representation of both of them in mice lacking Treg cells (Fig. 5B). In addition, pathway analysis showed an enrichment for terms related to anti-viral responses, cytokine secretion, and responses to IFNγ (Fig. S7A). GFAP+ astrocytes in the hippocampus exhibited a parallel reduction in arborization in Treg-deficient mice (Fig. 5C), suggesting widespread glial activation in the hippocampus.

**Figure 5.**
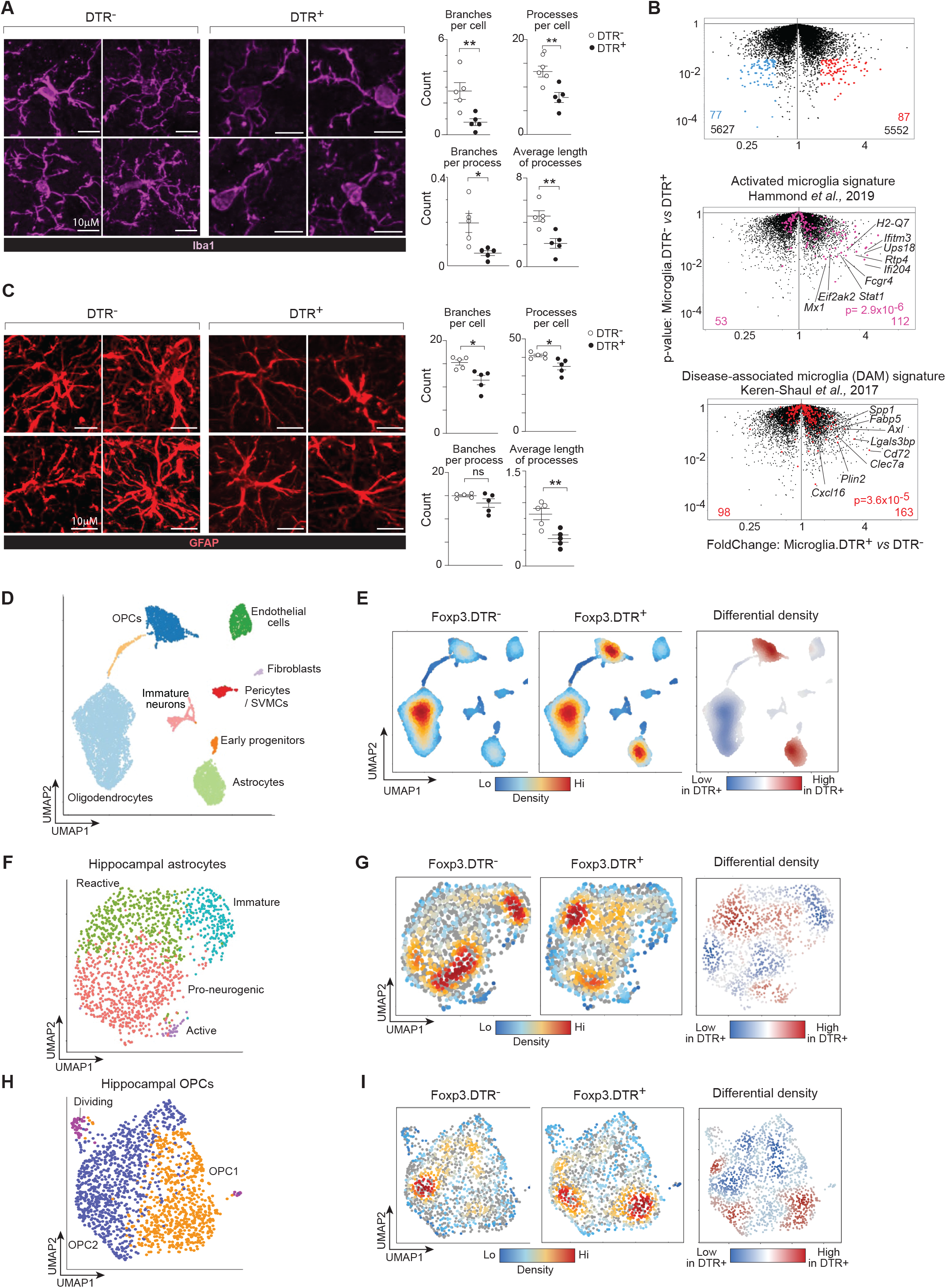
Treg-cell control of hippocampal glial cells. (A) Left: representative confocal images of DG microglia (Iba1+); right: summary quantification of classical morphological changes of activation, as described in the Methods section. n= 5 (B) “Activation upon demyelination” (*58*) (top) and “Disease-associated” (*57*) (bottom) microglia signatures are red-highlighted on a volcano plot comparing transcripts expressed in sorted hippocampal microglia from DT-treated DTR- versus DTR+ mice. Duplicate samples. (C) Same as panel A, but images of astrocytes (GFAP+). (D-I) scRNA-seq analyses of dissected hippocampal cells depleted of mature neurons (Thy1+) and microglia (CD45+). Duplicate samples. (D) UMAP representation. (E) Local cell densities on the UMAP space from DTR- (left) or DTR+ (center) mice. Density differential between the two genotypes (right). (F) UMAP representation of re-clustered astrocytes (light green) from panel D. (G) Same as panel E except specifically for the astrocyte cluster (light green) in panel D. (H-I) Same as panels F and G except the OPCs (blue) from panel D were re-clustered. OPCs, oligodendrocyte precursor cells; SVMCs, smooth vascular muscle cells; other abbreviations as per Fig. 3. Mean ± SEM, p-values as per Fig. 3.

For another view of the effects of Treg loss on hippocampal glia, we turned to scRNA-seq analysis. In duplicate, we dissected the hippocampus from DTR- and DTR+ mice, excluded mature neurons (Thy1+) and microglia (CD45+) so as not to obscure differences in rarer cell types, and performed scRNA-seq. After quality control, we retained a total of 10,234 cells with an average of 2,182 unique transcripts per cell. Dimensionality reduction, unsupervised clustering, and signature analysis revealed all of the major cell types known to participate in the neuronal stem-cell niche (*71*) (Fig. 5D) in both conditions (Fig. 5E). Although the same cell types were present in hippocampi from DTR- and DTR+ mice, their distribution differed in the presence vs absence of Treg cells, the hippocampus from DTR+ mice showing an enrichment for astrocytes and oligodendrocyte precursor cells (OPCs), the latter at the expense of oligodendrocytes (Fig. 5E).

Re-clustering of the astrocytes in isolation revealed four subtypes (Fig. 5F), the validity of which was confirmed by a heatmap of the twenty genes most differentially expressed by each subcluster (Fig. S7B). Given the identity of these genes, we have termed the four subtypes: immature, pro-neurogenic, reactive, and active, as per (*72, 73*). Upon punctual Treg-cell ablation, the hippocampal astrocyte landscape evolved from primarily immature and pro-neurogenic to reactive (Fig. 5G). An analogous procedure was applied to the OPC cluster, yielding two previously characterized subtypes: pro-myelinating (OPC2s) and pro-inflammatory (OPC1s) (*74*) (Fig. 5H, Fig. S7C, D). Loss of Treg cells provoked a switch to the pro-inflammatory phenotype (Fig. 5I).

## Treg cells regulate hippocampal neurogenesis

The hippocampus is one of the few brain regions where new neurons are generated in adult mice and humans (*75, 76*). Hippocampal neurogenesis is required for effective learning and memory formation throughout adulthood (*55, 77*). As illustrated in Fig. S8A, hippocampal pluripotent progenitors, i.e. radial-glia-like (RGL) cells, reside in the subgranular zone (SGZ) of the DG. As they differentiate into neurons, RGL cells proliferate and migrate axially, becoming intermediate progenitor cells for neurons (nIPC). nIPCs migrate vertically into the granular cell layer (GCL), where they are known as neuroblasts. Neuroblasts undergo a complex series of changes that resolves in either cell death or integration into the neuronal circuitry (*75, 78*).

Distilling and re-clustering neuronal cells from the scRNA-seq data of Fig. 5D, followed by marker analysis, revealed 5 clusters: early progenitors, proliferating IPCs, nIPCs, neuroblasts, and a few contaminating OPCs (Fig. S8B). Loss of Treg cells (in DTR+ mice) resulted in a very slight decrease in early progenitor cells and a small increase in nIPCs (Fig. S8B). In addition, the transcriptomes of the various subtypes were very similar in the presence and absence of Treg cells (Fig. S8C); although, suggestively, cells from the early-progenitor or nIPC clusters of DTR+ mice showed small reductions in several transcripts encoding proteins crucial for stem-cell identity (e.g. *Sox9, Hes1, Fezf2, Neurod1, Hmgn2*) (*55, 79–83*), differences that were no longer evident at the neuroblast stage.

To delve more deeply and specifically into potential changes undergone by early neural progenitors in the absence of Treg cells, we re-clustered them in isolation, yielding three discernable subtypes (Fig. S8D), populated by both scRNA-seq replicates (Fig. S8E). A heatmap of the most differentially expressed genes confirmed the clustering (Fig. S8F), and the gene identities prompted us to designate the subtypes “RGL cells,” “activated neural stem cells” (aNSCs), and “quiescent neural stem cells” (qNSCs). These designations were supported by heatmaps of transcripts encoding the key RGL-cell identifiers, *Gfap* and *Sox2* (Fig. S8G), and by overlaying on the UMAP published signatures of qNSCs and aNSCs (*84*) (Fig. S8H). RNA-velocity analysis, which uses the ratio of spliced to unspliced transcripts to predict the future transcriptional state of each cell, suggested with high confidence that RGL cells could differentiate into either qNSCs or aNSCs in the presence of Treg cells (Fig. S8I, left), as has been reported (*84*). Partition-based Graph Abstraction (PAGA) solidified this interpretation (Fig. S8I, left). Punctual ablation of Treg cells, resulted in loss of most of the velocity vectors (and, consequently, loss of directional confidence), suggesting a block of both of these differentiation routes (Fig. S8I, right), consistent with the fact that the biggest change in the RGL-cell compartment subsequent to Treg-cell ablation was an increase in the fraction of *Gfap+Sox2+* progenitor cells (Fig. S8J). In addition, Metascape analysis revealed the transcriptomes of hippocampal RGL cells from Treg-cell-depleted mice to be relatively enriched in transcripts associated with aerobic respiration (known to be a feature of NSC activation (*75, 85–87*)), stress, and apoptosis, for example. They were also relatively impoverished in transcripts associated with evolving RGL cells, such as radial-glia-like cell differentiation and gliagenesis (*88, 89*) (Fig S8K). Gene-Set-Enrichment analysis (GSEA) confirmed this reduction in pathways associated with neurogenesis and gliagenesis in RGL cells from mice lacking Treg cells (Fig. S8L). (Note also that these pathways did not come up when comparing 10 randomly re-shuffled datasets.)

## Treg cells protect RGL cells from IFNγ-induced death and mice from short- and long-term memory impairment

Seeing changes in neurogenesis after such a short Treg-cell depletion was not expected and begged independent confirmation. As an orthogonal approach, we performed quantitative stereological imaging of the DG after immunohistochemical staining of indicative markers. This approach side-stepped several drawbacks of scRNA-seq, e.g. tissue-dissociation artefacts, imperfect translation of mRNA abundance to protein abundance, low sequencing depth. Thus, we considered it a test of the hypothesis-generating scRNA-seq approach. The imaging approach also provided information not only on tissue organization in the presence vs absence of Treg cells but also on their numbers, morphology and location.

First, as illustrated in Fig. 6A, we divided the DG into the SGZ, which encompasses RGL cells, and the GCL, which hosts immature and mature neurons. There was no evidence of macro-morphological abnormality in the absence of Treg cells as we did not observe any differences in sections from DTR- vs DTR+ mice in either GCL volume (Fig. 6B) or SGZ area (Fig. 6C). There were also no significant differences in the number of immature (DCX+) neurons (Fig. 6D, Fig. S9A), and only a small increase in cellularity of the GCL zone (Fig. 6E). Focusing on the SGZ, we found significantly more RGL cells (SOX2+GFAP+) in DTR+ mice (Fig. 6F), consistent with the augmentation observed via scRNA-seq (Fig. S8J). An increase in RGL cells in the absence of more immature neurons prompted us to assess their state. There were more dead cells (Fractin+) in the SGZ of DTR+ mice (Fig. 6G), in the absence of an increase in proliferation [phosphorylated histone (pH)3+] (Fig. 6H). This observation was substantiated by scRNA-seq data demonstrating an increase in transcripts encoding pro-apoptotic BAX in the absence of Treg cells, with no change in transcripts specifying anti-apoptotic BCL2 (Fig. 6I) nor enrichment of proliferation signatures (Fig. S8K). Flow-cytometric analysis (gating scheme in Fig. S9B) also confirmed that RGL cells (CD45-SOX2+) from DTR+ mice were more frequently undergoing apoptosis (Annexin-V+ and cleaved-caspase-3+) in the absence than in the presence of Treg cells (Fig. 6J, Fig. S9C), and that this change was not compensated for by an increase in proliferation (Fig. 6K, Fig. S9D); more mature neurons showed the opposite dynamics (CD45-SOX2-) (Fig. 6E, Fig. S9C, D). Importantly, there was a significant negative correlation between the fraction of Treg cells in the meninges and the fraction of dead RGL cells in the hippocampus (Fig. 6L).

**Figure 6.**
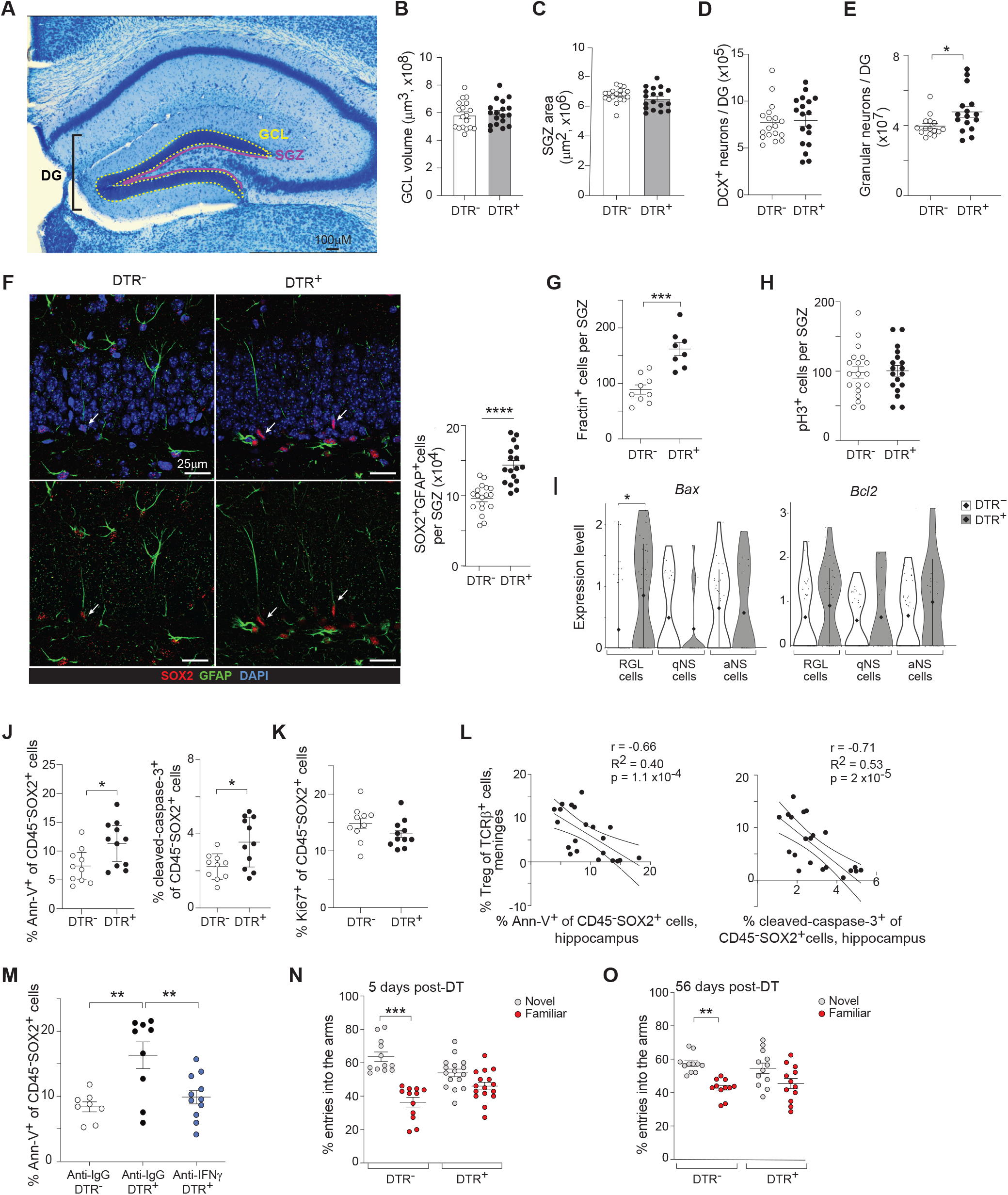
Treg-mediated protection of RGL cells from IFNγ-induced death and from short- and long-term memory impairments. (A-N) DTR+ and DTR- littermates (6 – 7wks old) were injected with DT as per Fig. 3A and analyzed on day 5 after the first injection. n=18 for panels B-F. (A) Representative image of Nissl staining of the hippocampus. Relevant DG anatomical structures: SGZ (solid magenta line), GCL (dotted yellow line). (B) GCL volume. (C) SGZ area. (D) Histological quantification of immature (DCX+) neurons. (E) Histological quantification of granular neurons. (F) Left and center: representative confocal images of RGL cells (arrows) from DTR- and DTR+ littermates. All markers are shown in the upper panels, while DAPI staining has been removed in the lower panels. Right: summary quantification of RGL cells per SGZ. (G) Histological quantification of Fractin+ cells per SGZ. n≥8 (H) Histological quantification of pH3+ cells per SGZ. n≥18 (I) Violin plot of *Bax* and *Bcl2* transcription expression from the clusters delimited in Fig S8D. (J) Flow-cytometric quantification of the frequency of RGL cells (CD45-SOX2+) expressing death markers: Annexin-V+ (left) and cleaved-caspase-3+ (right). n≥10 (K) Percentage of Ki-67+CD45-SOX2+ cells. n=11 (L) Correlation between the meningeal Treg frequency and the frequency of death markers on the RGL cells (CD45-SOX2+): Annexin-V+ (left) and cleaved-caspase-3+ (right). n=21 (M) Flow-cytometric analysis of Annexin-V+ RGL (CD45-SOX2+) cells from DT-treated DTR+ and DTR- mice treated with an anti-IFNγ or isotype-control mAb, as detailed in the Methods section. n≥8 (N) Frequency of an individual mouse’s entries into the familiar (red dots) versus novel (grey dots) arms of the Y-maze 5 days after the initial DT injection. n≥12 (O) Same as panel M, except the readout was 56 days after the initial DT injection. n≥11 DG, dentate gyrus; GCL, granular cell layer; SGZ, subgranular zone; IFN, interferon; other abbreviations as per Fig. 6. Mean ± SEM, p-values as per Fig. 3.

Because, as illustrated in Fig. 4I, punctual ablation of Treg cells resulted in a strong enrichment of IFNγ-signaling pathways in the hippocampus, we wondered whether this cytokine was involved in the neuronal-cell death provoked by loss of Treg cells. IFNγ has been reported to have a concentration-dependent effect on RGL cells, mostly in *in vitro* settings: at low concentrations, it promotes differentiation of RGL cells into neurons, while at higher concentrations it is toxic (*13, 29, 87, 90–95*). Hippocampal RGL cells expressed both subunits of the IFNγ receptor (Fig. S9E). Furthermore, IFNγ-response genes were specifically enriched in the RGL-cell cluster of Fig. S8D (Fig. S9F). Under the DT-injection protocol illustrated in Figure 3A, we co-administered an anti-IFNγ mAb or an irrelevant anti-IgG control mAb 8 hours before the first DT injection and then every other day until day 5. Neutralization of IFNγ was indeed able to reduce the fraction of dead RGL cells in the hippocampus to the levels of Treg-containing mice (Fig. 6M), without affecting death of non-RGL cells (Fig. S9G). Despite the ability of anti-IFNγ to curb death, the CD4+ and CD8+ effector T cell populations still expanded (Fig. S9H), placing the latter event upstream of IFNγ production. To determine whether IFNγ could have a direct impact on RGL-cell survival, we generated neurospheres from the hippocampus of 3-10-day-old mice (*96, 97*) and cultured them with versus without recombinant IFNγ (rIFNγ) according to two protocols (Fig. S9I). When rIFNγ was added at the initiation of neurosphere formation, there was a dose-dependent reduction in live RGL cells (Fig. S9J) as well as the appearance of dead-cell blebs (Fig. S9K) a week later, likely signaling a reduction in the initial pool of hippocampal RGL cells. Parallel results were obtained when preformed neurospheres were treated with rIFNγ (Fig. S9J).

Lastly, to determine whether the inhibition of RGL-cell differentiation provoked by loss of Treg cells had a functional impact on the hippocampus, we used a Y-maze test to evaluate memory formation. Spatial-reference memory, which is dependent on the hippocampus (*55*), was tested by placing a mouse in the Y-maze with one arm closed off during a training period. After a resting period, the mouse should remember which arm it already explored and visit it less frequently than the unexplored arm. Whereas DTR- mice did remember the familiar arm and entered the novel arm more often, DTR+ mice did not show a significant preference, indicating an inability to remember what they had previously explored (Fig. 6N). Surprisingly, this defect was maintained for at least 8wks after Treg-cell depletion, demonstrating that the functional damage to the hippocampus was not just short-term (Fig. 6O).

## Treg cells control IFNγ levels by depriving T and NK effector cells of IL-2

What mechanism(s) underlay Treg control of T- and NK-cell activities in the meninges and brain parenchyma, in particular their production of IFNγ? We previously reported that punctual depletion of local Treg cells unleashes proliferation of, and IFNγ production by, NK and T cells in the pancreatic islets. And that a major mechanism by which Treg cells operate in this context is depriving effector cells of IL-2 because Tregs’ high-level display of CD25, the high-affinity-conferring chain of the IL-2 receptor, renders them powerful competitors for this critical growth and survival factor (*98*). Treg cells are known to commonly rely on this “cytokine-sink” property to suppress immune responses (*99*). We hypothesized that a similar scenario might be playing out in the meninges upon abrupt Treg depletion. In support of this notion, near-total ablation of CD25+ Treg cells using an anti-CD25 mAb –with a much milder effect on total Treg-cell numbers – induced a meningeal IFNγ response that mimicked surprisingly closely the one provoked by punctual depletion of the entire Treg compartment in DT-treated *Foxp3.Dtr+.Gfp+* mice (Fig. S5C-E).

IL-2 binds with high affinity to a trimeric receptor consisting of IL-2Rα (CD25), IL-2Rβ (CD122), and the cytokine common γ chain (γc, CD132), and with low affinity to a dimeric receptor composed of the latter two chains; both receptors signal through a STAT5-directed transcriptional program (*100*). Staining meningeal immunocytes from wild-type mice revealed frequent phosphorylated STAT5 (pSTAT5) expression by CD4+ Tconv, CD8+ T, NK, and B cells, suggestive of IL-2 sensing, but not by γδT cells or MFs. Littermates punctually depleted of Treg cells (as per Fig. 3A) showed significantly increased pSTAT5-positivity for only CD4+ Tconv, CD8+ T, and NK cells, pointing to an IL-2 boost (Fig. 7A). This interpretation was supported by the induction of a published IL-2-response signature in NK cells and CD4+ Tconv cells 24 hours after a single injection of DT into *Foxp3.Dtr+.Gfp+* mice (Fig. S10A), less impressively for the T cells likely due to their slower activation kinetics (*101, 102*).

**Figure 7.**
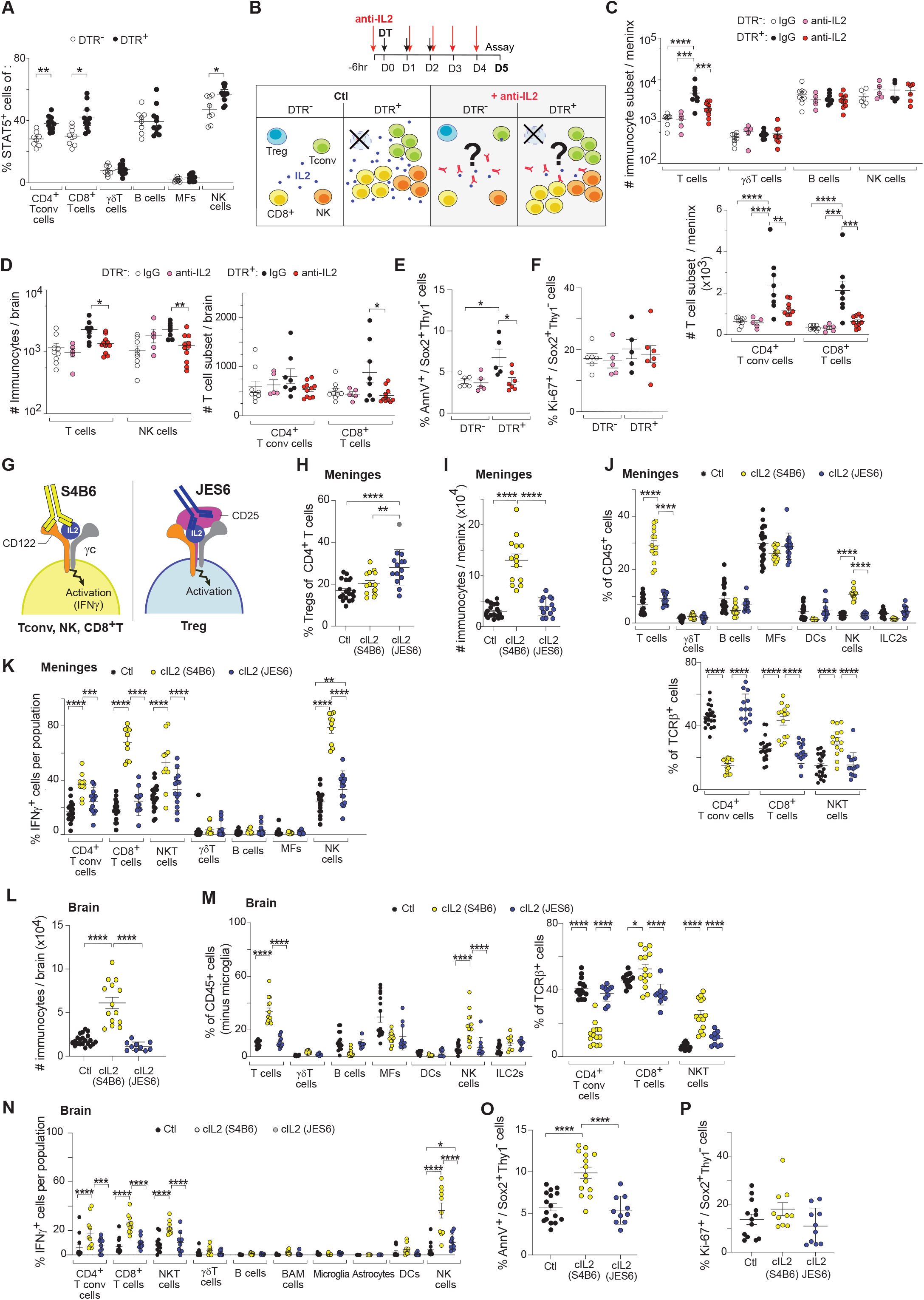
Duramater IL-2 availability is fine-tuned by Treg cells. (A) Frequency of pSTAT5+ select immunocyte populations of mice depleted of Treg or not. n ≥ 8 (B-F) Flow-cytometric quantification of DTR+ and DTR- male littermates (6-7wks old) ip-injected with DT and anti-IL2 or isotype control mAb, as detailed in the Methods section. n ≥ 5 (B) Anti-IL2 blocking protocol and schematic of the different conditions. (C) Numbers of meningeal select immunocyte populations (top) and T cell subsets (bottom). (D) Same as panel C except for the brain parenchyma. (E) Frequency of Annexin-V+ hippocampal RGL cells. (F) Frequency of Ki-67+ hippocampal RGL cells. (G-P) Flow-cytometric analysis of the duramater and brain parenchyma of B6 WT littermate males (6 – 7wks old) after ip-injection of PBS (black), JES6-containing complex IL2 (cIL2) (gray) or S4B6-containing cIL2 (white). n ≥ 14 (G) Schematic representation of the both cIL2 and their target cells (H) Percentage of meningeal Tregs. (I) Total meningeal immunocyte counts. (J) Percentage of meningeal select immunocyte populations (left) and T cell subsets (right). (K) Percentage of IFNγ+ immunocyte populations. n ≥ 10 (L) Total brain immunocyte counts. n ≥ 10 (M) Same as in panel I except for the brain. n ≥ 10 (N) Same as in panel J except for the brain. n ≥ 10 (O) Frequency of Annexin-V+ hippocampal RGL cells. n ≥ 10 (P) Frequency of Ki-67+ RGL cells. n≥10 Mean ± SEM, p-values as per Fig.3. For simplicity, only relevant statistical differences are shown.

If Treg-cell control of IL-2 availability is indeed important for guarding homeostasis of the meningeal immune system, co-incidental Treg depletion and neutralization of IL-2 should prevent the usual expansion of and IFNγ production by meningeal T and NK cell populations. Neutralization of IL-2 in the presence of Treg cells, as per the protocol depicted in Fig. 7B, did not significantly change the numbers of total meningeal Treg cells (Fig. S10B), total immunocytes (Fig. S10C), or any lymphocyte subset examined (Fig. 7C). In contrast, neutralizing this cytokine in littermates concomitantly depleted of Treg cells strongly reduced expansion of the total, CD4+ Tconv and CD8+ T cell populations (Fig. 7C). Similarly, neutralization of IL-2 in mice hosting Treg cells had little effect on immunocyte accumulation within the brain parenchyma, while parallel neutralization in Treg-deficient littermates inhibited the typical expansion of CD8+ T and NK cells at that site (Fig. S10D, E and Fig. 7D). IL-2 neutralization also reduced the death of hippocampal RGL cells characteristically provoked by Treg-cell depletion (Fig. 7E) without impacting their cell-cycle entry (Fig. 7F), thereby linking expansion and parenchymal infiltration of T and NK populations with IFNγ-mediated RGL-cell death. In addition to the lack of an anti-IL-2 effect in Treg-containing mice, the specificity of these observations is argued by the lack of splenic T and NK population expansion in anti-IL-2-treated, Treg-deficient mice (Fig. S10F-H).

Additional support for the notion that Treg control of IL-2 availability was key to maintaining meningeal immune-system homeostasis came from IL-2-supplementation experiments: does specifically increasing the IL-2 available to T and NK cells surmount their control by Treg cells and promote their accumulation? Injection of IL-2/anti-IL-2 complexes containing the antibody clone S4B6 has been shown in several systems to highly preferentially agonize the low-affinity IL-2R dimer expressed by CD4+ Tconv, CD8+ T and NK cells but not the high-affinity IL-2R trimer expressed primarily by Treg cells (owing to the cytokine conformation locked in by this particular mAb) (*98, 100, 103*); in contrast, injection of IL-2/anti-IL-2 complexes containing the antibody clone JES6 better agonizes CD25-containing IL-2Rs, thereby preferentially expanding Treg populations (see schema in Fig. 7G). Indeed, the meningeal Treg pool was not expanded 3 days after injection of IL-2/S4B6 complexes but was increased with parallel injection of IL-2/JES6 complexes (Fig. 7H). In contrast, there was a large increase in total meningeal immunocytes after injection of S4B6-containing, but not JES6-containing, complexes (Fig 7I), by far the largest fold-changes being those of T and NK cells (Fig. 7J). There was also a striking increase in the fractions of T and NK cells expressing IFNγ (Fig 7K). The brain parenchyma showed parallel changes with IL-2/S4B6 vs IL-2/JES6 complex injection: many more total immunocytes (Fig. 7L), above all T and NK cells (Fig. 7M) and an expansion of IFNγ+-expressing T and NK cells with S4B6-containing complexes (Fig. 7N). Moreover, there was more hippocampal RGL-cell death with injection of S4B6-containing complexes, with no changes in the fraction of dividing cells (Fig 7O, P).

Another major mechanism of Treg-cell suppression of other immunocytes *in vivo* is secretion of anti-inflammatory cytokines, in particular IL-10 (*18, 104, 105*), which is important for tissue homeostasis in several contexts, notably in the colon (*18, 106*). IL-10 has also been implicated in neuronal survival and control of neuroinflammation (*24*). We flow-cytometrically compared the immunocyte compartments of 6-week-old *Foxp3-Cre.Il10wt/wt* and *Foxp3-Cre.Il10fl/fl* littermates, i.e. mice devoid of IL-10 production specifically by Treg cells. We found no significant differences in the numbers of any immunocyte compartment examined (Fig. S10I, J).

On the basis of this series of experiments, we propose the following mechanistic scenario (as depicted in Fig. S11): In unperturbed wild-type mice, meningeal Treg cells keep the activation, proliferation, IFNγ production and parenchymal infiltration of T and NK cell effectors in check by out-competing them for limiting amounts of the critical growth/survival factor, IL-2. Upon punctual loss of Treg cells, the T and NK populations become activated, rapidly expand, and become potent IFNγ producers. They also develop the capacity to invade the brain parenchyma, where they activate glial cells, especially in the hippocampus, and promote death of hippocampal RGL cells, likely both directly and indirectly, thereby compromising their differentiation. Consequently, short- and long-term spatial-reference degrades

## DISCUSSION

A series of studies over the past several years has revealed the richness of the meningeal immune system in mice, as well as the important roles it can play in brain responses to environmental and physiological challenges such as aging, injury, or infection (*34, 107*). Most of these explorations concentrated on the cellular and molecular impacts of various myeloid (MF) or lymphoid (T or B cell) effector populations. Our study, in contrast, focused on a largely uncharacterized population of meningeal Treg cells and the regulatory mechanisms they can mobilize to safeguard brain homeostasis at steady-state.

Besides controlling meningeal immunocyte responses, including impeding their access to the brain parenchyma, Treg cells exerted widespread influences on the hippocampus. Upon acute loss of Treg cells, the entire hippocampal region switched to a reactive, pro-inflammatory state, without any evidence of the compensatory dampening mechanisms reported to arise in more progressive brain insults such as with experimental autoimmune encephalomyelitis or in the early stages of neurodegenerative diseases (*59, 108–110*). RGL cells suffered the most dramatic changes, Treg depletion blocking their differentiation and inducing their death. To some extent, these effects may result from the profound changes in glial-cell states. For effective neurogenesis to occur, the various glia types must cooperate to clear the niche and, together, must provide appropriate signals to RGL cells to support their differentiation into newborn neurons (*61, 62, 71, 111*). As discussed below, the RGL-cell changes almost certainly reflected direct influences of IFNγ as well. Regardless, Treg-cell depletion appeared to leave a “scar” in the hippocampus, engendering long-term functional defects in memory formation. Notably, a pattern of hippocampal changes very similar to that induced by punctual ablation of Treg cells – pan-glia activation plus RGL cell death coupled with accumulation – was recently observed in several human neurodegenerative diseases (*76*).

How do meningeal Treg cells supervise the RGL-cell niche? Vis-à-vis their lymphoid-organ counterparts, meningeal Tregs preferentially express a number of transcripts that encode proteins capable of influencing neurogenesis and brain function more generally: *Il10*, *Wnt3*, *Hpgds, Psap* or *Ptgs1,* amongst others (*24–27, 112*). And there exist several precedents for meningeally produced cytokines diffusing into the brain and having profound impacts on cognitive behavior (*8–11, 16, 17*). But our data highlight a more indirect scenario: meningeal Treg cells kept local IFNγ-producing T and NK cells in check. Punctual ablation of Treg cells led to rapid production of IFNγ in the meninges, which ultimately resulted in RGL-cell death and sustained defects in neurogenesis within the DG. Thus, meningeal Tregs would not be expected to recognize antigens expressed by NSCs. Rather, they would be most likely to share specificities with the meningeal (and eventually brain) CD4+ and CD8+ effector T cells they are counteracting. Such a scenario is highly reminiscent of the recent proposal by Medzhitov and Iwasaki of introspective, self-reactive “Tx” cells that are critical for tissular homeostasis and are counterbalanced by tissue-Tregs (*113*)

We were struck by the similar cast of immunocytes that infiltrated the hippocampus (or the brain more generally) in the Treg-depletion model on one hand and with aging or neurodegeneration in mice and humans on the other. Like the loss of Treg cells, aging is associated with increased levels of IFNγ (*114*), coupled with T and NK cell invasion of the brain parenchyma (*115–117*). Relatedly, T cells have been increasingly implicated as key pathogenic agents in human and mouse neurodegenerative diseases such as Parkinson’s disease, Lewis body dementia, and AD (*118–122*). For example, in a mouse model of AD, T cells – especially CD8+ T cells – producing IFNγ infiltrated the brain parenchyma, in particular the hippocampus, and were a major driver of disease progression (*120*). Intriguingly, in AD patients, Treg cells were one of the immunocyte subsets collected from the cerebrospinal fluid that was most changed in comparison with those from healthy controls (*123*). But how and why these pathogenic, clonally expanded T cells access the brain parenchyma has remained a mystery. A malfunction of meningeal Treg cells – perhaps due to aging (*124*) or reflecting some imbalance in the various meningeal Treg subtypes, or due to a break in the meninges-parenchyma communication axis (*34*) – culd potentially be causal or exacerbating.

## Limitations

The fact that many of our experiments entailed organismal depletion of Foxp3+ cells raises the question of whether the effects we observed primarily reflect a local or systemic loss of Treg cells. Several observations argue that depletion of Treg cells in the meninges was responsible for at least some of the defects manifest in mice rendered Treg-deficient via our standard protocol. First, we performed assays at a quite short time after the depletion protocol in order to mitigate systemic effects. Second, injection of an anti-CD25 mAb into the intracisternal magna, a means to remove Treg cells that is less inflammatory (*45*) and is meninges-preferential (*33*), provoked meningeal changes very similar to those observed upon DT-mediated Treg depletion. Third, very early time-course analyses revealed a meningeal response to IFNγ that was earlier and encompassed more cell types than did the splenic response (as systemic-response indicator). And, lastly, perturbations of the brain parenchyma were localized to specific functional regions, notably the hippocampus, and were absent from others, e.g. the cerebellum. Nonetheless, we cannot completely rule out systemic effects from the Treg-depletion protocol, via circulation of either cells or mediators. Nor can we completely rule out a dominant input from the leptomeninges or choroid plexus, although this seems unlikely given that they host 15x and 49x fewer Treg cells than the dura. Regardless, the salient finding is that Treg cells safeguard brain homeostasis in healthy mice in a multi-pronged fashion.

## MATERIALS AND METHODS

### Experimental model and subject details

#### Mice

*Foxp3IRES-GFP* (termed B6.*Foxp3Gfp* here) mice (*125*) were obtained from Dr. V. Kuchroo (Brigham and Women’s Hospital, Boston, MA), while *Foxp3DTR* (termed *B6.FoxP3Dtr* here) mice (*47*) came from Dr. A. Rudensky (Memorial Sloan Kettering Cancer Center, NY, NY). 6wk- to 7wk-old male mice were used for all experiments unless otherwise specified. For RNA-seq experiments, 6wk- or 15wk-old male mice were used. Mice were housed in our specific-pathogen-free facility at Harvard Medical School (HMS). For the time-course experiment, 3wk-, 52wk-, and 72wk-old C57BL/6J (B6) mice were purchased from the Jackson Laboratory and acclimated in our facility for at least 3 wks. Balb/cJ, NOD/ShiLtJ and C3H/HeJ mice were purchased at 3 wks of age from Jackson Laboratory and acclimated in our facility for at least 3 wks. All experiments were conducted under a protocol (#IS00001257) approved by HMS’s Institutional Animal Care and Use Committee.

#### Various mouse treatments

For all the experiments, mice were intracardially perfused with at least 35 mL of cold phosphate-buffered saline (PBS) before harvesting tissues. For ablating Treg cells, B6.*Foxp3Dtr+* mice and B6.*Foxp3Dtr-* littermates of the designated ages were intraperitoneally (ip) injected with diphtheria toxin (DT) (Sigma) at 20 ng/g body weight for 3 consecutive days. For intravascular labeling of immunocytes, we followed the protocol reported in (*53*). Briefly, the mice were anesthetized and injected with 25 μL of mouse anti-CD45 mAb (BioLegend) intracardially (left ventricle). After 3 minutes, 500 μL of blood was extracted to confirm the blood-immunocyte labelling. Lastly, we perfused the mouse with 35 mL of cold PBS and harvested the tissues of interest. For IFNγ neutralization, we injected rat Ultra-LEAF™ purified anti-mouse IFNγ mAb (BioLegend) or the corresponding rat IgG1κ isotype-control Ab (BioLegend). We ip-injected 250 μg per mouse every 48 hours, starting 8 hours prior to the first DT injection. For anti IL2 neutralization, we injected ultra-LEAF purified anti-mouse IL2 mab (JES6-1A12, Biolegend) or the corresponding rat IgG2ak isotype-control Ab (Biolegend). We ip- injected 100ug per mouse every 24 hours, starting 6 hours prior to the first DT injection. For IL-2 treatments, Il-2-anti-Il-2 complexes were prepared as previously described for the different anti-Il-2 mAb clones, JES6-1A12 (*126*) or S4B6 (*98*). Shortly, for the clone that binds to the high affinity IL-2 receptor (CD25): we incubated 0.5ugr mouse Il-2 (PepreoTech) to 5ugr of an IL-2 mAb (JES6-1A12, Biolegend) for 30 minutes on ice, resuspend it in 100uL of cold sterile PBS followed by intraperitoneal injection. For the clone that binds to the lower affinity IL-2 receptor (CD122): we incubated 5ugr mouse Il-2 (PepreoTech) to 50ugr of an IL-2 mAb (S4B6; BD or BioXcell) for 30 minutes on ice, resuspend it in 100 uL of cold sterile PBS followed by intraperitoneal injection.

#### Intracisterna-magna injections

Injections into the cisterna magna were used to deliver anti-CD25 mAb or an isotype control mAb into the subarachnoid space in order to deplete meningeal Treg cells. The procedure was performed as previously described (*51*). Mice were first anaesthetized with isoflurane (Thermo Fisher Scientific) (4%) in an induction chamber and treated with carprofen (20 mg kg–1, subcutaneous) and buprenorphine (0.1 mg kg–1, subcutaneous) immediately before the surgery. Lubrication of corneas was maintained using Puralube (Dechra), and anaesthesia by isoflurane was maintained through a nose cone during the surgery. Animals were transferred to a stereotaxic frame, and the skin of the head was shaved and aseptically prepared by swabbing betadine followed by ethanol (three times each). The cranium was exposed by making a surgical anterior–posterior incision with a scalpel blade, and the subcutaneous tissue and muscles of the neck were gently separated to access the dura mater of the cisterna magna. 2.5 μg of Ultra-LEAF™ purified mouse anti-CD25 mAb (BioLegend, clone PC61) or the recommended rat IgG1λ isotype control mAb (BioLegend) were injected into the subarachnoid space using a 30-gauge 0.5-inch needle mounted on a 25- μl Hamilton syringe. After injection, muscles were re-aligned, and the incision was closed using wound clips (Autoclip, 7 mm) and tissue adhesive (Vetbond, 3M). After surgery, animals were placed in a cage containing a warming pad and monitored for 1 hour. Additional doses of carprofen (every 24 h) were administered for 72 h after surgery, and wounds were monitored for adequate healing.

### Isolation of leukocytes from lymphoid and non-lymphoid tissues

#### Dura mater, leptomeninges, and choroid plexus dissection and digestion

All of the following steps were performed on ice and/or on cold blocks in order to preserve tissue quality. After perfusion, to isolate the dura mater, we removed the bottom part of the skull, exposing the undamaged brain parenchyma. We carefully extracted the brain and stored the skullcap with the adhered meninges in cold RPMI-1640 medium (Thermo Fisher Scientific) with 2% fetal bovine serum (FBS) until all dissections were done. Each meninge was digested in 2 mL of a digestion cocktail made of: RPMI, 2% FBS, 0.1 μg/mL in DNAse I (SIGMA), and 0.5 mg/mL collagenase P (ROCHE). Each skullcap was placed into a small tissue-culture dish (FALCON, REF 353001) filled with cold RPMI/FBS and, under a dissecting microscope, peeled the dura mater from the skull, and then put in a 2 mL microtube (AXYGEN, ref MCT-200-C-S) filled with digestion buffer. The samples were incubated with shaking at room temperature (RT) for 45 min. Next, we prepared 50 mL conical tubes with 100 μm cell strainers (FALCON, ref 352360) on top. We used a 1mL pipette tip pre-coated in RPMI/FBS to reduce stickiness. We also pre-wet the filter with 2 mL of RPMI/FBS to avoid filter stickiness and to have some volume at the bottom of the tube. We pipetted up and down at least 15 times to totally disaggregate the meninges. Then, we filtered it and washed twice to remove any cells that might be left behind. The filter was washed with 15 mL of RPMI/FBS. The tubes were spun at 520x*g* for 6 minutes, the supernatant aspirated, and the pellet resuspended in the desired volume of RPMI/FBS.

#### Skull bone-marrow dissection

Once the meninges were removed, we took the skullcap, placed it in a 1.5 mL Eppendorf tube with 1 mL of RPMI/FBS and finely chopped it. We filtered the sample through a 70 μm cell strainer to remove big bone pieces, washed it with 10 mL of RPMI/FBS, and spun it down at 520x*g* for 5 minutes, followed by red-blood-cell lysis (Gibco) and washing with 10 mL of RPMI/FBS.

#### Spleen preps

Immunocytes were obtained by mechanical disruption followed by red-blood-cell lysis, were filtered with a 40 μm strainer, and were washed with 10 mL of RPMIf solution.

#### Brain dissection and digestion

To extract various regions of the brain, we placed each brain [stored in cold HBSS (Hanks’ Balanced Salt solution) (Gibco)] into a tissue-culture dish on top of a cold block in order to reduce cell death and better maintain its integrity. Regions of interest were dissected under a microscope, carefully removing the choroid plexus to avoid contamination, and were placed in a 2 mL microtube. To obtain a single-cell suspension, we used the Milteny neural dissociation kit (T) (ref 130-093-231), following the manufacturer’s instructions, but with a few modifications. Instead of using GentleMACS, we used manual mechanical dissociation. After a 15min incubation at 37°C under slow continuous rotation, we transferred the “Enzyme Mix 2”. Then, in a 37°C water bath, we gently pipetted up and down 15 times per tube, repeating this process for a total of 20 minutes, followed by filtering and washing per the manufacturer’s instructions. For the whole-brain digestion, we first minced the brain to allow for a better digestion and then followed the same protocol as described above.

#### Neurospheres

Hippocampal neurospheres were isolated using a modified protocol from Zhang, Y., Hu, W. (*127*). Briefly, hippocampi were harvested from male neonatal pups (5-10 days post birth). To isolate the RGL cells, hippocampi were washed with ice cold HBSS then carefully mechanically digested using a razor blade or scapple. Hippocampi were then placed in a 15 mL conical tube. Then they were added to 3 ml of prewarmed (37°C) dispase [1U/mL] digestion medium (StemCell Tech 07923). The samples were incubated at 37°C for 12 min while inverting the tube every 3 minutes. Digested material was spun down (200 rcf for 5 min at 4°C) and supernatant discarded. 3 ml of prewarmed collagenase type IV (1mg/mL) digestion medium (StemCell Tech 07909) was added to cell pellet. Samples incubated the at 37°C for 10 min while inverting every 3 min. Next, each sample was centrifuged (200 rcf for 5 minutes at 4C) and supernatant discarded. Cells were washed twice with HBSS. In a 50 ml tube, tissue was resuspended in 10 ml prewarmed (37°C) 0.05% Trypsin-EDTA diluted in HBSS. and incubated for 3-5 min while shaking in a 37°C water bath, immediately followed by adding cold 13 ml digestion neutralizing medium (DMEM/F12 [10% FBS]) into Trypsin-EDTA solution. Solution was spun at 200 rcf for 5 min at 4C and supernatant was removed. 1 ml of digestion neutralizing solution was added and the sample resuspended by pipetting up and down. Then, cells were filtered through a 70um strainers and washed with neutralizing medium twice. Finally, the cells were pelleted at 200 rcf for 5 min and resuspended in 1 ml of Neurosphere Proliferation Medium consisting of NeuroCult Basal Medium (StemCell Tech 05700) containing 5 mL NeuroCult Proliferation Supplement (StemCell Tech 05701), 100 ul EGF [10 ug/mL], 50 ul bFGF [10 ug/mL], 50 ul Heparin Solution (StemCell Tech 07980), and 500 ul Antibiotic/Antimycotic (100X). Cells were counted using a hemocytometer. Cells were plated at 5.15 x 104 live cells per well in 1 mL of Neurosphere Proliferation Media.

RGL cells were stimulated with IFNγ in two different ways. For the pretreatment with IFNγ. The moment cells were plated IFNγ was administered into the Neurosphere Proliferation Media at the desired concentration (1ng/mL or 10ng/mL). Neurospheres were collected 7 days post treatment. In other experiments, to assess IFNγ treatment on established neutrosphere cultures, neurospheres were cultured without IFNγ for 7 days. On day 7, IFNγ was administer every 48 hours and neurospheres collected 72 hours after the first IFNγ administration.

To process the neurospheres at the desired timepoint, they were spun down (200 rcf,1 minute), and the pellet digested in accutase for 15 minutes. Next, we added twice the volume of DMEM and centrifuge it (210 rcf, 5 minutes 4C), the supernatant aspirated, and pellet resuspended in the desire volume of RPMI/FBS.

#### Flow cytometry

The following antibodies were used for flow cytometric staining: anti-CD45, -CD8a, -CD4, -TCRγδ, -CD64, -CD11b, -CD11c, -IA/IE, -CD25, -CD19, -CD45R, -CD24, -CD43, -IgD, -IgM, -CD279, -CD278, -CD62L, -IL-17A, -IFNγ, -IL4, -IL5, -TNFα, -Thy1.1, -Tbet, - NK1.1, -Ly6G, -CX3CR1, -CD38, -Sox2, -Annexin V, -CD69, -CD138, -Streptavidin BV605, -CD3, -Ly6C, -CD137, -F4/80, and -CD86, all from BioLegend; anti-Foxp3, - RORγt, -ST2, and -IL13 from eBioscience; anti-TCRβ, -Ki-67, -ID3, and -CD185 from BD (Becton, Dickinson and Company); anti-cleaved-caspase-3, -TCF1 from Cell Signaling; and -S1P1/EDG-1 from R&D. Surface staining was performed for 30 minutes at 4°C, and viability was assessed using Zombie UVTM Fixable Viability Dye, Zombie Yellow Fixable Viability Dye, or Zombie NIR Fixable Viability Dye (BioLegend) as per the manufacturer’s instructions. Intracellular staining was performed using eBioscience’s Foxp3/Transcription Factor Staining Buffer Set. For *ex-vivo* intracellular cytokine staining, single-cell suspensions were stimulated for 3.5 hours at 37°C with 50 ng/mL phorbol 12-myristate 13-acetate (PMA) and 1 mM ionomycin (both from Sigma-Aldrich) in the presence of protein transport inhibitor cocktail (eBioscience) in complete RPMI medium supplemented with 10% FBS. Cells were acquired with a FACSymphony flow cytometer (BD Biosciences). Data were analyzed using FlowJo software (Tree Star).

### Imaging

#### Immunohistochemistry of whole-mount dura mater

Mice were perfused with HBSS and then decapitated. The skullcap containing the dura mater was removed from the head and drop-fixed overnight in a 1:1 mixture of BD ICC Fixation Buffer and HBSS at 4 °C. Skullcaps were then washed in HBSS and stored in HBSS + 0.1% NaN3 at 4 °C. The dura was carefully removed from the skull cap using forceps and permeabilized/blocked in 0.3% Triton-X100 + 20% donkey serum + 5 μg/mL anti-mouse Fc Block in HBSS for 1 hour at RT. Dura maters were then stained with a cocktail of primary antibodies diluted in 0.3% Triton-X100 + 20% donkey serum in HBSS overnight at 4°C. Staining solution was removed, and tissues were washed 3 times for 1 hour at RT in 0.3% Triton-X100 in HBSS. In some cases, tissues were then stained with species-specific secondary Abs diluted in 0.3% Triton-X100 + 20% donkey serum in HBSS overnight at 4°C followed by 3 washes, then replacement with HBSS. Dura whole-mounts were then flattened onto a microscope slide, allowed to dry at RT, and cover-slipped with ProLong Diamond Antifade solution (Thermo Fisher scientific). Slides were allowed to dry overnight at RT before imaging.

#### Confocal imaging of dura whole-mounts

Tissues were imaged on a Zeiss LSM 710 or Zeiss LSM 880 confocal microscope with Zen Black software using a 10x air objective or 25x, 40x, or 63x oil immersion objectives. Images from the same experiment were acquired with identical digital bandpass, laser, gain, and pinhole settings, and the pinhole aperture was set to the same value across imaging tracks. Tile-scan images were collected with a 10% overlap and stitched in Zen Black.

#### Confocal image processing and manual cell quantification

Images were processed in Fiji v2.1.0. For each experiment, the minimum and maximum display range was adjusted manually for one image and then batch-applied to the rest of the images. Individual images are generally displayed as a single confocal plane or a stack of several adjacent planes. Tile-scan images are displayed as maximum-intensity projections. Treg cell counts on tile-scan images were obtained by manually annotating individual Foxp3-GFP+ cells, and then overlaying the annotation onto a map where the borders of the dural sinuses were demarcated. Cells were then designated as “sinus-” or “non-sinus-” associated.

#### Generation of cell-density heatmaps

Maximum projections of tile-scan images were preprocessed through a Fiji ImageJ macro to set reproducible minimum and maximum display intensities per channel. Individual channels were then thresholded to create binary images. The alpha-smooth muscle actin (αSMA) channel of each image was registered to a standardized template of the transverse and sagittal dural sinuses using a point-based geometric transformation in MatLab. Registration points for the αSMA channel were then applied to the remaining channels of each image. For a given channel [e.g. GFP (Foxp3)], registered images from multiple mice were aggregated by calculating the arithmetic mean-intensity per pixel. Pixels were then binned by rescaling the averaged image matrix to create superpixels. A pseudocolor heatmap was created using these superpixel values, representing the relative frequency with which signal from a specific cell type [e.g. GFP (Foxp3)-expressing Treg cells] was found at a given X-Y coordinate across multiple mice. Pseudocolor heatmaps were then overlaid on an aggregate, binarized image of the αSMA+ dural sinuses to visualize cell-type location.

### Immunohistochemistry of the brain parenchyma

#### Tissue collection

For standard brain histology, the mice were perfused with at least 35 mL of cold PBS, then perfused again with 35 mL of 4% paraformaldehyde (PFA) (Electron Microscopy Science). For immunohistochemistry (IHC) of cFos+ cells, brain astrocytes (GFAP+), and microglia (Iba+), were collected in 15 mL Falcon tubes and placed overnight (ON) at 4 °C. The next morning, they were layered over 10% sucrose at 4°C. Once brains sank to the bottom of the tube (6-8 hours), they were layered over 20% sucrose at 4°C and the process repeated; then layered over 30% sucrose solution and the process repeated. The next day, brains were placed on a mold with Tissue-Tek® O.C.T. Compound (SAKURA), frozen, and stored at -80°C until processing. Five or more non-consecutive sections of 20 μm thickness were stained in order to sample areas throughout the hippocampus and other brain regions. Sections were preserved on the slides at -30°C until use.

For staining, the sections were thawed at RT for 2min and then washed twice at RT with PBS (five minutes each) to rehydrate them. The samples were permeabilized and blocked in PBS-Tx (PBS, 0.25% Triton X-100) supplemented with 1:100 Fc block, 2% bovine serum albumin (BSA) (Sigma-Aldrich), and 5% normal donkey serum (blocking solution) (Jackson ImmunoResearch Laboratories) for 2hrs at RT. They were then incubated in PBS-TX plus blocking solution with the primary antibody (cFos (Cell Signaling) 1:500, CD3 (Abcam) 1:100, Iba1 (Abcam) 1:100, GFAP (Abcam) 1:500) overnight at 4°C. The samples were rinsed and washed three times (10 minutes each) in PBS-Tx solution at RT, and then incubated with the secondary antibody, a donkey against the primary antibody species (1:500) for 2 hrs in PBS-Tx and blocking solution at RT. The following steps were performed in the dark to avoid bleaching of the labelled Abs: another round of 3 washes with PBS-Tx; an extra wash of 10min with PBS; a 5min incubation at RT with 4’,6-diamino-2-phenylindole (DAPI, Sigma-Aldrich); and three 5min washes with PBS. The sample was cover-slipped with ProLong Diamond Antifade solution, and the slides allowed to dry overnight at RT before imaging. Images were acquired with a Nikon Ti2 inverted microscope, employing either a widefield and Plan Apo 20x air objective or a spinning-disk confocal and a Plan Fluor 40X oil objective. Data analysis and quantification were performed using FIJI software (ImageJ1 version 2.9.0).

### Imaging for neurogenesis

#### Histology

Mice were perfused as above for standard IHC, stored in 4% PFA/PBS and shipped to Spain. Serial coronal brain sections (50 µm thick) were obtained from the left hemisphere on a Leica VT1000S vibratome and individually collected into a 96-well plate filled with PBS. Plates were kept at 4 °C until further analysis.

#### Immunohistochemistry

Each series of sections entailed systematic sampling of the hippocampus in the rostro-caudal axis (8-9 hippocampal sections per animal, 50 µm thick sections 400 µm apart, with one random series chosen for each staining). Slices were initially preincubated in PBS with 5% Triton X-100 and 0.1% BSA (PBST-BSA) for single- or double-staining. Cell nuclei were counter-stained with 4’,6-diamino-2-phenylindole (DAPI, Sigma-Aldrich). Another randomly chosen series was used for Nissl staining in order to determine the total volume of the granular cell layer (GCL) of the DG and the area of the subgranular zone (SGZ). Primary antibodies: anti-SOX2, goat anti-sex determining region Y-box 2 (1:200, R&D Systems, af2018); GFAP, rabbit anti-glial fibrillary acidic protein (1:1000, Abcam, Ab7260); pH3, rabbit anti-phospho-histone H3 (1:500, Millipore, 06-570); DCX, goat anti-doublecortin (1:500, Santa Cruz, sc-8066); CLR, rabbit anti-calretinin (1:3000, Swant, 76994); Fractin, rabbit anti-Fractin (N-terminal fragment of actin) (1:1000, Millipore, AB3150) Abs were incubated with agitation in PBT-BSA for 1 hr at RT and 72 hrs at 4°C. Secondary Abs (Donkey anti-goat alexafluor 594, Donkey anti-rabbit AlexaFluor 594, Donkey anti-rabbit AlexaFluor 488; Invitrogen; 1:1000) were incubated in PBT-BSA for 1hr at RT and 24 hrs at 4C.

#### Stereology

For the hippocampal volume estimations, the Cavalieri method was used essentially, as described previously (*128, 128*). Briefly, we imaged (Leica, Thunder microscope) the hippocampal formation from Nissl-stained samples. We delimited regions of interest and measured both SGZ length and GC area (ImageJ). These measurements were extrapolated using Cavalieri’s method [the sum of the lengths or areas of each slice multiplied by the distance (d) between each slice (d = 400 μm)].

Fractin+ and pH3+ cells, representing dead and proliferating cells, respectively, were counted in the series of vibratome sections under a STELLARIS 8 Confocal Microscope and an optical fluorescence microscope (Leica DMI 6000 B), respectively. The total number of labelled cells was estimated by the optical fractionator method, multiplying the cell count by 8 (i.e. 1/8th of every hippocampal section of an animal was counted; thus the sampling fraction was 8). For death, only Fractin+ cells showing DNA condensates in the DAPI signal were counted. For proliferation, only cells staining for pH3, located in the SGZ, and showing mitotic morphology were counted.

For SGZ SOX2+/GFAP+ cells, 7 confocal stacks per mouse, corresponding to the SGZ and GCL of the DG, were obtained. Dissectors were randomly positioned in rostro-caudal sections of the GCL of the DG visibly containing the SGZ. All physical dissectors were obtained with confocal microscopy (Leica TCS SP5, oil immersion 40× objective + 2.4 digital zoom). We identified and counted [using Image J (Fiji 1.53)] only cells that met these criteria: 1) SOX2+/GFAP+ staining; 2) cell body positioned in the SGZ; and 3) RGL-cell morphology with a long process across the GCL. The total number of early progenitors per mouse was estimated by multiplying the cell density by the SGZ area of each animal (estimated using the Cavalieri method).

Doublecortin- and/or calretinin-expressing cells (DCX+/CLR+ and DCX+/CLR- cells) were counted using Image J (Fiji 1.46) on 6 physical dissectors for each animal, obtained via confocal microscopy (Leica TCS SP5, oil immersion 63X objective + 2.46 digital zoom). Dissectors were randomly positioned in rostro-caudal sections of the GCL of the DG visibly containing the SGZ. The density of DCX+/CLR+, DCX+/CLR-, and DCX-/CLR+ cells was calculated per each dissector. The total number of the various populations of neural precursors along the hippocampus was calculated as above for SOX2+/GFAP+ cells.

For the total number of granular cells in the DG, we took four confocal stacks (63x, + 7.5 digital zoom). Every cell inside the dissector was a granular cell. We estimated the density for each stack and the mean per animal. The total number of granular cells was calculated by multiplying the mean density by the volume of the GCL.

### RNA-seq methodology and analyses

#### Gene signatures

We used the following gene signatures: pan-tissue-Treg signature from (*19*); Th1-like-Treg signature and IFN-responsive-Treg signature from (*42*); Tfr-like-Treg signature from (*43*); quiescent-Treg signature from (*36*); OPC1 and OPC2 signatures from (*74*), activated and quiescent neuronal-stem-cell signatures (*84*); disease-associated-microglia signature from (*57*), activated microglia from (*58*); MCAO and LPS signatures from (*54*); EAE-reactive-astrocyte signature from (*59*). The inflammatory signatures, IFN-related signatures and neurogenesis-related signatures are from the Gene Ontology (GO) database (https://www.informatics.jax.org/vocab/gene_ontology) and GSEA (MsigDB) database (https://www.gsea-msigdb.org/gsea/msigdb/index.jsp). More specifically: Inflammatory response (GO 0006954), Antigen processing and presentation (GO 0019882), Cellular response to IFNγ (GSEA-GOBP), Cellular response to type II interferon (GO 0071346), IFN gamma response (GO 003434), Type II interferon signaling (GO), IFNγ mediated signaling pathway (GSEA-GOBP), Positive regulation of inflammatory response (GO 0050729), Neuroinflammatory response (GO 0150076), Response to virus (GO 0009615), Interferon alpha and beta signaling (GSAE-Reactome), Response to interferon-alpha GO 0035455, Response to interferon-alpha (GO 0035455), Type I interferon-mediated signaling pathway (GO 0060337), Type I interferon response (GO 0034340), Neurogenesis (GO 0022008), Neuron differentiation (GO 0030182), Neuron development (GO 0048666), Central nervous system neuron development (GO 0021954), Regulation of neurogenesis (GO 0050767), myelin sheath (GO 0043209). IFNγ response (GSEA/MsigDB, “HALLMARK_INTERFERON_GAMMA_RESPONSE”).

#### Population-level RNA-seq library construction, sequencing and data processing

Samples were double-sorted using a FACSAria or FACS Astrios: 600-1000 cells from each population were collected into 5 μl TCL buffer (Qiagen) containing 1% beta-mercaptoethanol (β-ME) (Sigma) in DNA LoBind tubes (Eppendorf). Library construction, sequencing and data processing was performed according to ImmGen’s standard operating procedures (https://www.immgen.org/img/Protocols/ImmGenULI_RNAseq_methods.pdf).

The DESeq2 package (Bioconductor) was used to normalize the raw read-counts via the median of ratios method and then converted to the GCT and CLS formats. Samples with less than 1 million uniquely mapped reads were excluded from normalization. Multiplot Studio (developed by Scott Davis, Mathis/Benoist lab) was used for visualization of transcriptional data. Transcripts were further filtered for minimal expression (>10 reads) and coefficient of variation cutoffs (<0.60).

RNA was isolated from whole tissues using TRIzol (Invitrogen) following the manufacturer’s instructions. 2ng of RNA in 5 μL of TCL containing 1% β-ME in DNA LoBind tubes were used for library construction, sequencing, and data processing, as described above.

#### scRNA-seq and analysis

For the scRNA-seq dataset on meningeal Treg cells, we merged data from two independent experiments (using the Harmony algorithm). We sorted CD4+ T cells from meninges of 15wk- and 36wk-old B6 males. Due to the low number of cells in each experiment, samples were pooled and tagged (hashed) with DNA-coded anti-CD45/MHCI Abs (BioLegend) to differentiate them from other samples in the same sequencing lane. Cells were encapsulated using the Chromium Single Cell 5′ v3 and V(D)J platform (10x Genomics). Data were processed using the standard CellRanger pipeline (10x Genomics). Single-cell data were analyzed following the Seurat v4 pipeline. Hashtag oligonucleotide (HTO) counts were obtained using the CITE-seq-Count package. HTOs were assigned to cells using the HTODemux function, and doublets and negative cells were eliminated from analysis. Low-quality cells and doublets were also excluded from the analysis according to the criteria in Table S2. Clusters of Treg cells were identified on the basis of expression of Treg marker genes (e.g., *Foxp3*, *Il2ra*, *Ctla4*, and *Ikzf2*), while clusters of non-Treg cells were removed from downstream analysis using the SubsetData function. Signature scoring was done with Seurat’s AddModuleScore function. Visualization of expression density of transcripts or signature scores was performed using the Nebulosa package. Differential-density analysis was performed using the MASS package, employing the kde2d() function. We used the scVelo package (*129, 130*) for velocity analysis.

The *Tcr*-seq library was processed using the CellRanger vdj pipeline (v4.0 and v6.0), employing default parameters. This pipeline was used for the assignment of V, D and J genes, CDR3 regions and sequences. Repeated clonotypes were defined as cells sharing the VDJC segments of both *Tcra* and *Tcrb* with identical *Cdr3* sequences at the nucleotide level.

For the hippocampus scRNA-seq dataset, CD45-Thy1-cells were sorted by flow cytometry and encapsulated using the Chromium Single Cell 30 v3 platform. Two DTR- and two DTR+ samples were individually encapsulated. Samples were processed, hashed, and sequenced together to minimize batch effects.

#### Pathway analysis

Pathways available from the GO database (https://www.informatics.jax.org/vocab/gene_ontology/), GSEA (MsigDB) database and/or Metascape online tool were used (*131*).

### Behavioral Studies

#### Novel Y-maze

Behavioral testing was conducted by the Mouse Behavior Core at Harvard Medical School. Only male Foxp3DTR littermates were used for the behavioral studies. Mice were injected with DT at 6wks-old. The Novel Y-maze test is conducted in 3 phases: habituation, inter-trial interval (ITI), and test. During the habituation phase, one arm of the maze (either left or right) is randomly blocked off by a plastic insert, allowing mice to explore a random assortment of the other two arms of the Y-maze. The mice all began the maze from the same starting arm. After the habituation phase, each mouse was placed back into the holding cage for the ITI. The maze was cleaned with Windex and the blockade was removed prior to the test phase. During the Test phase, the mice were placed back at the starting arm and allowed to explore all 3 arms uninhibited. After each mouse had completed the test, the cages were thoroughly cleaned with Windex to remove any odors, hair, or feces.

For the young mice (6wks-old), we followed the standard protocol: a 3-min habituation phase and a 3-min test phase, spaced by a 2-min ITI. For older mice (15wks-old), we modified the times as per (*15*): each mouse had 10 mins habituation, 60 mins of ITI, and 10 mins of test. Mouse tracking and data analysis was performed with EthoVision XT 14 software.

### Statistical analyses

All statistical analyses were performed using GraphPad Prism software. If not stated otherwise, data are presented as mean ± SEM. Statistical significance was calculated by an unpaired Student’s *t* test (two-tailed), except for multiple comparisons (Figs. 1F and 7L), when the statistical significance was calculated using a one-way ANOVA. As indicated in the figure legend, p < 0.05 was considered significant. P-values for gene-signature enrichment or impoverishment between a pairwise comparison via volcano plots or FC/FC plots were determined using the χ2 test. \**p* < 0.05, \*\**p* < 0.01, \*\*\**p* < 0.001, ****p < 0.0001.

#### Data and materials availability

New data generated in this paper have deposited in the Gene Expression Omnibus (GEO) database under the accession number GSE234317.

## ACKNOWLEDGMENTS

We gratefully acknowledge: M.R. Kasal for manuscript editing; B. Hanna, A. Muñoz-Rojas, V. Piekarsa, K. Hattori and A. Ortiz-Lopez for experimental help and discussions; P.V. Anekal and the HMS MiCRoN Core for imaging help and guidance with image analyses; J. Lee, I. Magill, and the Broad Genomics Platform for RNA-seq; L. Yang and B. Vijaykumar for computational help; C. Laplace for graphics; B. Caldarone and the HMS Mouse Behavior Core; and the HMS Immunology Flow Cytometry Core.

## AUTHOR CONTRIBUTIONS

MMR and DM conceptualized the study MMR, ECR, AJW, TJ, FAPR and QR performed experiments and analyzed data RJ, IMC, BS, JLT, CB and DM provided funding RJ, IMC, BS, JLT and DM provided supervision MMR and DM drafted the manuscript MMR, ECR, AJW, TJ, FAPR, QR, RJ, IMC, CB, BS, JLT and DM reviewed manuscript MMR and DM edited writing

## List of Supplementary Materials

Figures S1 to S11

Table S1 to S2

References 132-135

## SUPPLEMENTARY FIGURE LEGENDS

**Figure S1.**
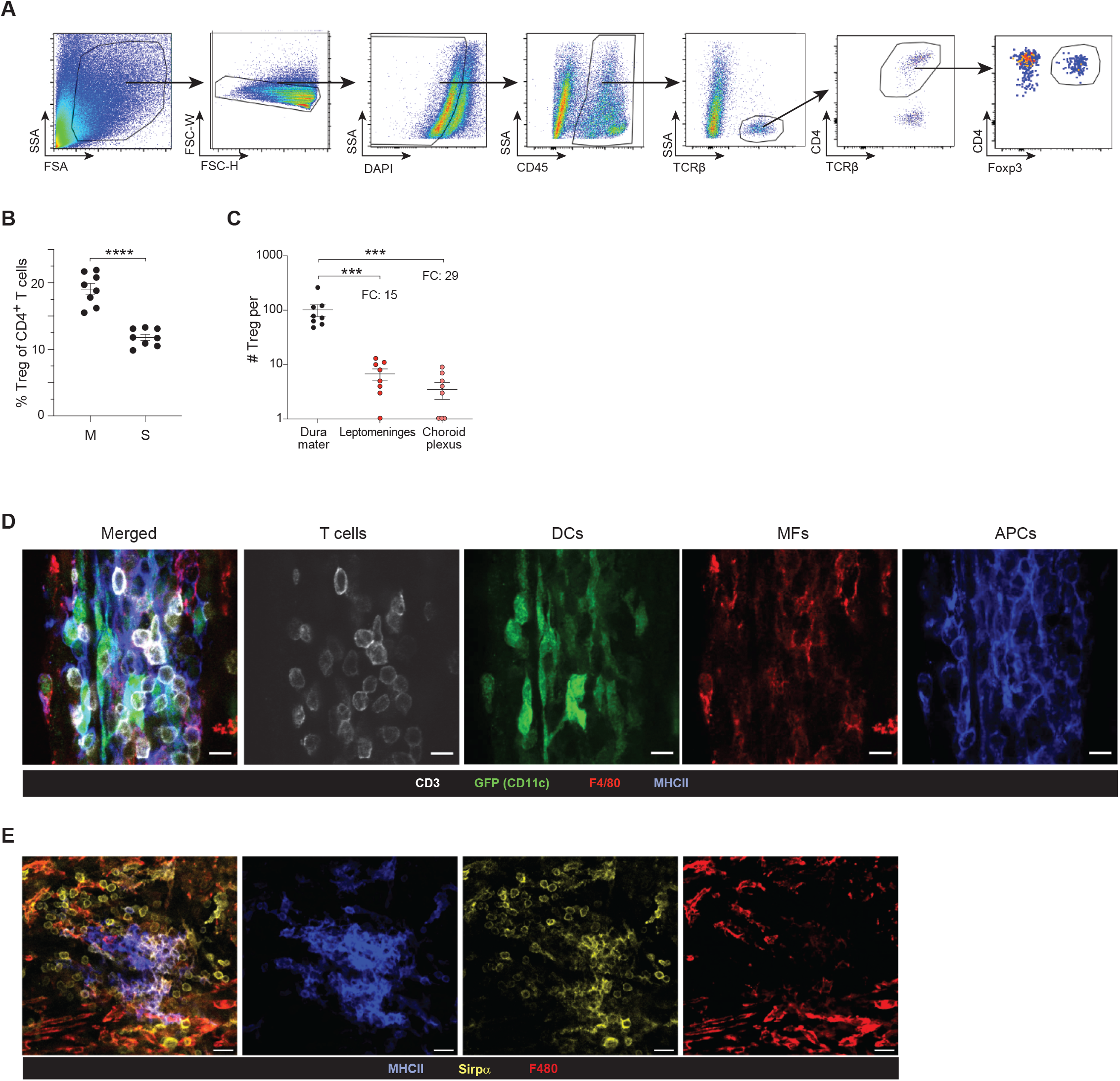
Supporting flow-cytometric and histological data. (A) Gating strategy for the flow-cytometric identification of meningeal Treg cells. (B) Treg-cell frequency in the meninges and in the spleen. n=8 (C) Quantification of Tregs in the different brain border regions. n=8 (D-E) Representative confocal microscopy of MF- and cDC2-defining markers, stained as indicated. DC, dendritic cell; MF, macrophage, APC, antigen-presenting cell; M, meninges; S, spleen

**Figure S2.**
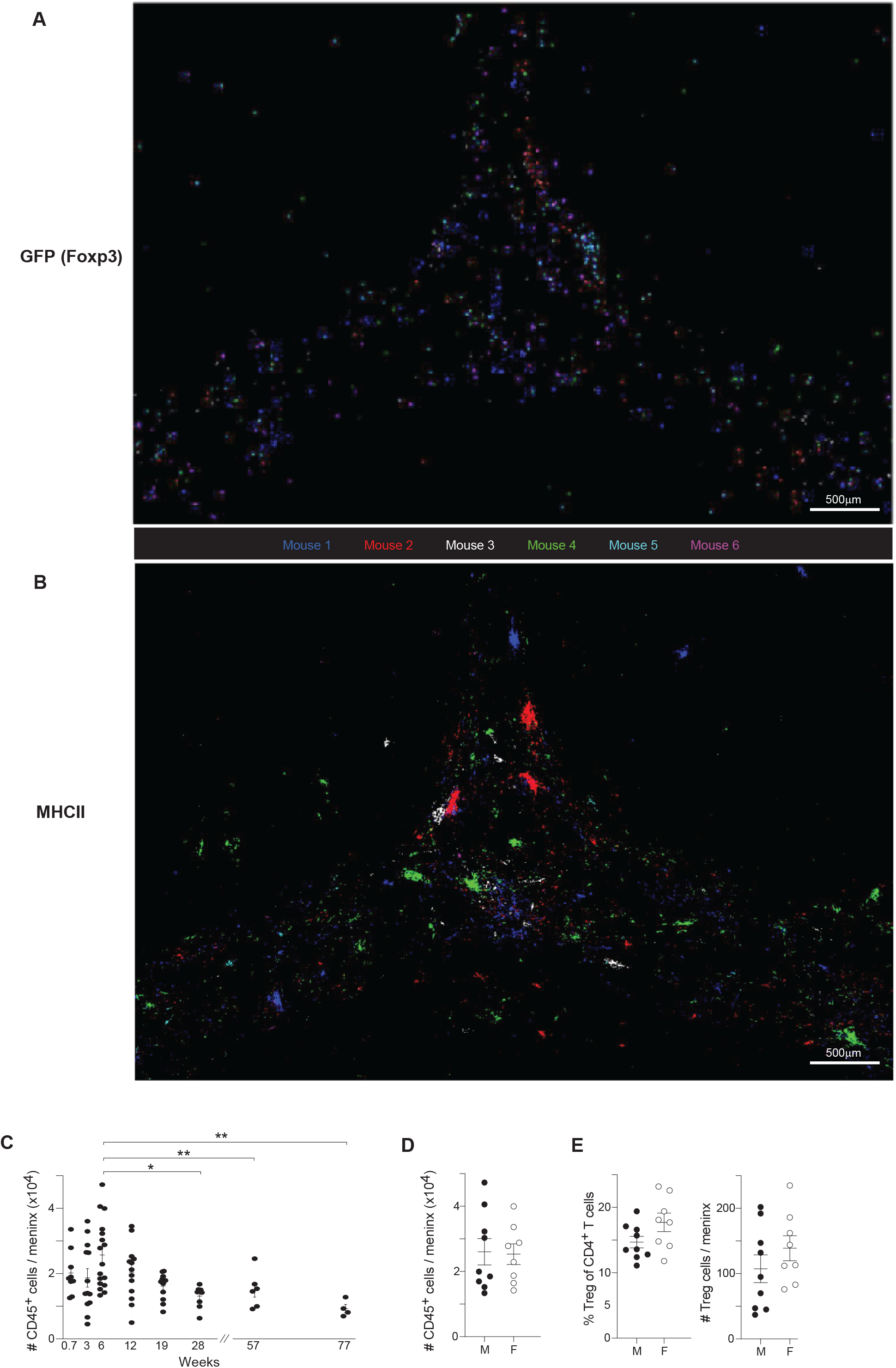
Additional data on Treg-cell localization and on responses to physiological variations in the meninges. (A-B) Related to Figure 1E, registered tile-scan images of multiple duras, depicting GFP positive (A) and MHCII+ (B) cells for individual mice, used to calculate densities. (C) Quantification of total immunocyte numbers in meninges of B6 mice of various ages. (D-E) Summary flow cytometric-data comparing 6wk-old male and female mice: total immunocyte counts (D); Treg-cell frequency (E, left) and numbers (E, right). Mean ± SEM. p-values as per Fig. 1.

**Figure S3.**
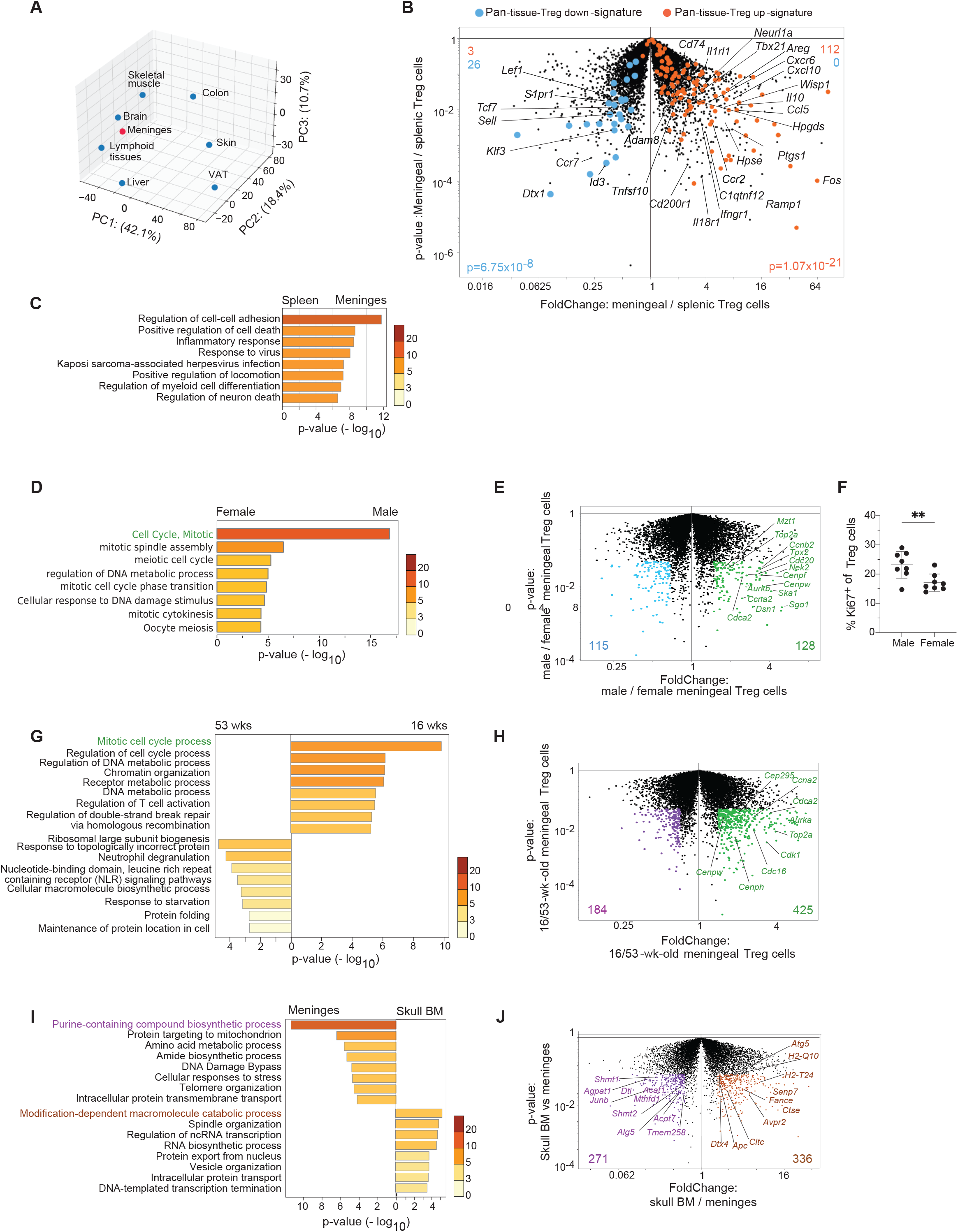
Additional characterization of meningeal Treg cells. (A) Principal component analysis (PCA) of population-level RNA-seq on double sorted Treg cells. Transcripts differentially expressed between each tissue Treg population and its corresponding lymphoid-tissue control (FC>2 and FDR<0.1) were calculated using the edgeR package. Data-sets from visceral adipose tissue (VAT), skin, liver, skeletal muscle, colon and EAE brain are from (*23, 132–134*). PCs 1, 2 and 3 with their proportions of explained variance are plotted. (B) Volcano plot comparing gene expression of meningeal and splenic Treg cells from 15wk-old male mice. A published pan-tissue Treg signature (*19*) is highlighted. Orange: up-regulated; blue: down-regulated. Triplicate samples. (C) Pathway-enrichment analysis via Metascape (*131*) of the transcripts differentially expressed in meningeal Treg cells (FC >1.5, p<0.05). (D-F) Transcriptomic differences between the meningeal Treg cells of male and female mice. (D) Pathway-enrichment analysis via Metascape (*131*) of the transcripts differentially expressed by meningeal Treg cells from male and female mice. (E) Volcano plot highlighting proliferation-related transcripts. Duplicate samples. (F) Flow-cytometric analysis of Ki-67+ meningeal Treg cells in the two sexes. n=8 (G-H) Transcriptomic differences between the meningeal Treg cells of 16wk- and 53wk-old male mice. (G) Pathway-enrichment analysis via Metascape (*131*) of the transcripts differentially expressed by meningeal Treg cells of the two ages. (H) Volcano plot highlighting proliferation-related transcripts. Duplicate samples. (I-J) Transcriptomic differences between Treg cells from the skull bone-marrow and meninges. (I) Pathway-enrichment via Metascape (*131*) analysis of transcripts differentially expressed in Treg cells from skull BM vs the meninges. (J) Volcano plot with key genes from enriched pathways from panel (H). Triplicate samples for skull BM Treg cells, duplicate samples for meningeal Treg cells. EAE, experimental autoimmune encephalitis. Mean ± SEM. p-values as per Fig. 1.

**Figure S4.**
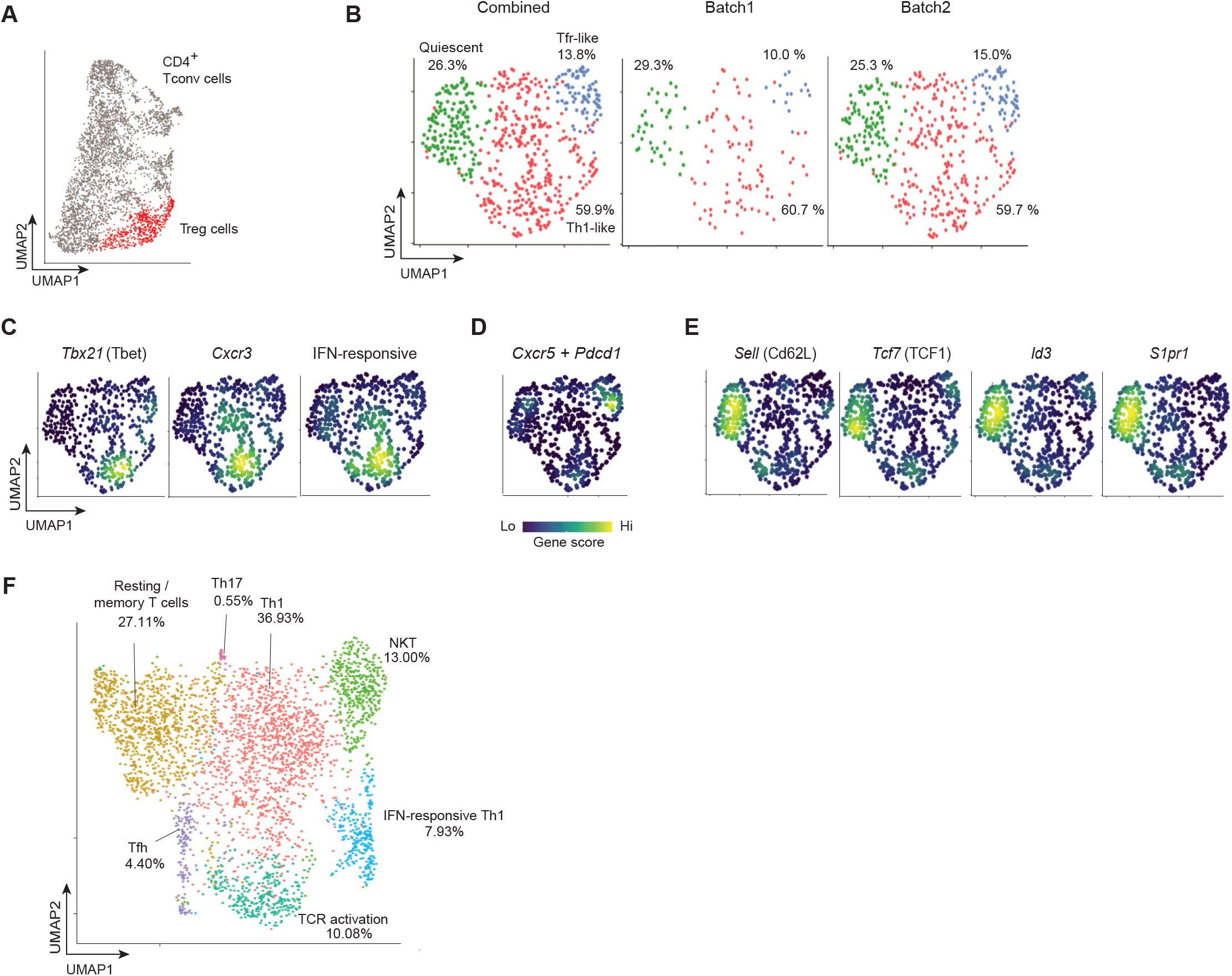
Additional characterization of meningeal Treg. (A) Merged UMAP representation of sorted meningeal CD4+ T cells from two independent experimental batches. (B) UMAPs of reclustered Treg cells (red-highlighted in panel A) illustrating the contributions of the two independent samples. Percentages refer to the fraction each subtype makes up the total of that sample. (C-E) Density plots of key genes and signatures for each of the meningeal Treg clusters from Fig 2C. (C) Th1-like Treg-cell markers: *Tbx21*, *Cxcr3*, IFN-responsive Treg-cell signature (*42*). (D) Tfr-like Treg-cell markers: *Cxcr5* + *Pdcd1* combined gene score. (E) Quiescent Treg cluster markers: *Sell, Tcf7, id3, S1pr1*. (F) UMAP of non-Treg CD4+ T cells. Treg cells were removed from the combined data of panel A (leaving the grey cells). Cell-type assignments were made by most differentially expressed genes. Percentages indicate the fractional representation of each cluster.

**Figure S5.**
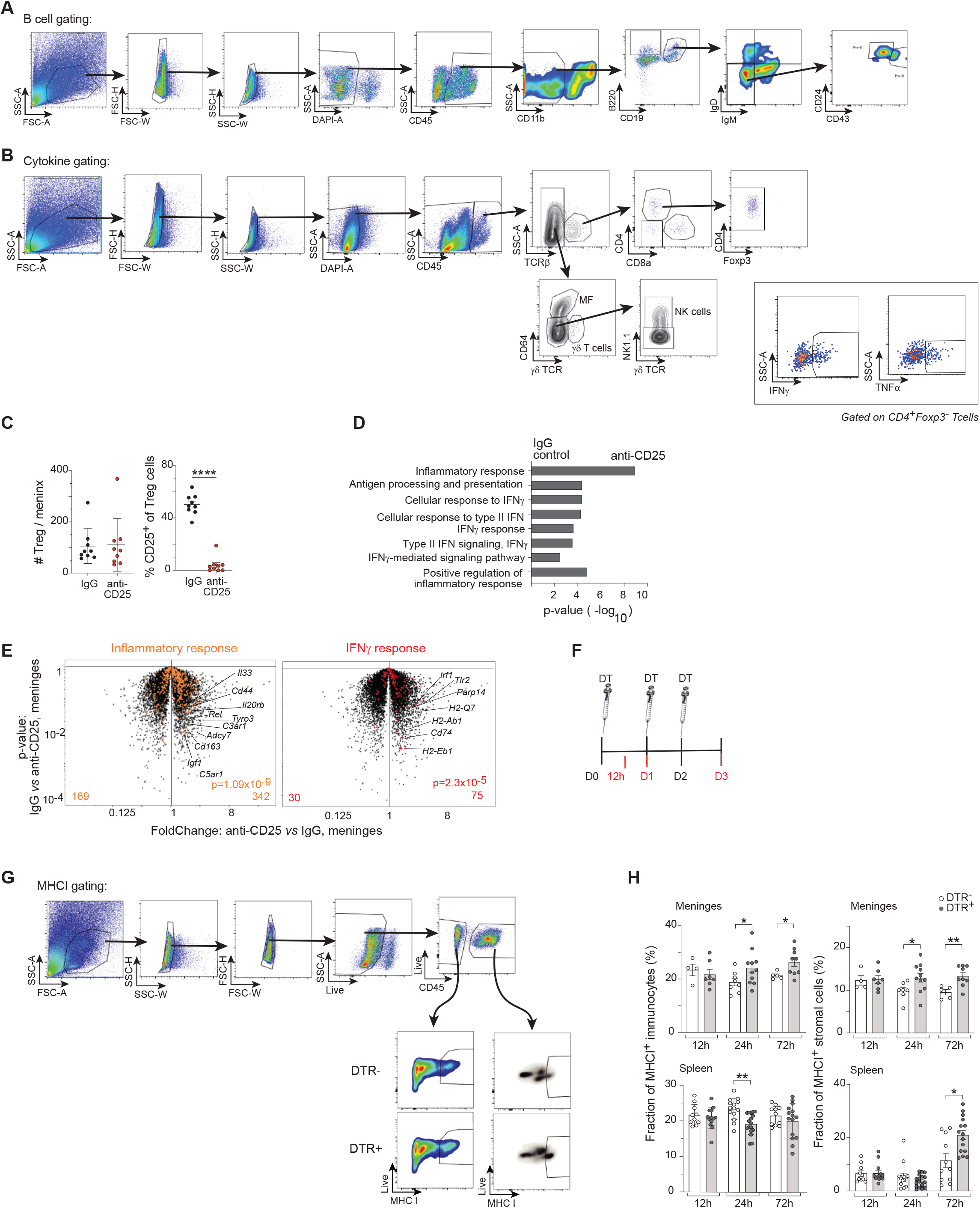
Demonstration of a classical meningeal, rather than a systemic, response to Treg depletion. (A) Gating strategy for the flow-cytometric identification of meningeal B cells. (B) Gating strategy for the flow-cytometric identification of cytokines in different meningeal immunocyte populations. (C-E) Analysis of the meninges three days after injection of anti-CD25 or an isotype-control Ab into the intracisternal magna. (C) Flow-cytometric quantification of the number of meningeal Treg cells (left) and their fraction of CD25+ cells (right). n≥8 (D) Pathway-enrichment analysis of differentially expressed genes (FC>1.5; p-value <0.05) from E of selected gene signatures from GO dataset and GSEA/MSigDB dataset related to: inflammatory response, IFNγ response, and type I IFN response. (E) “Inflammatory response” and “IFNγ response” from the Gene Ontology (GO) database are highlighted on a volcano plot of transcripts from meninges of mice treated with αCD25 or control IgG Ab. Triplicate and duplicate samples. (F-I) Time-course analysis early after DT injection into DTR+ vs DTR- mice. (F) Experimental scheme. (G) Gating strategy for the flow-cytometric identification of MHCI+ cells. (H) Flow-cytometric analysis of MHCI+ immunocytes (left panels) or stroma (right panels) isolated from the meninges (top row) or spleen (bottom row). n≥4 Mean ± SEM. p-values as per Fig. 1.

**Figure S6.**
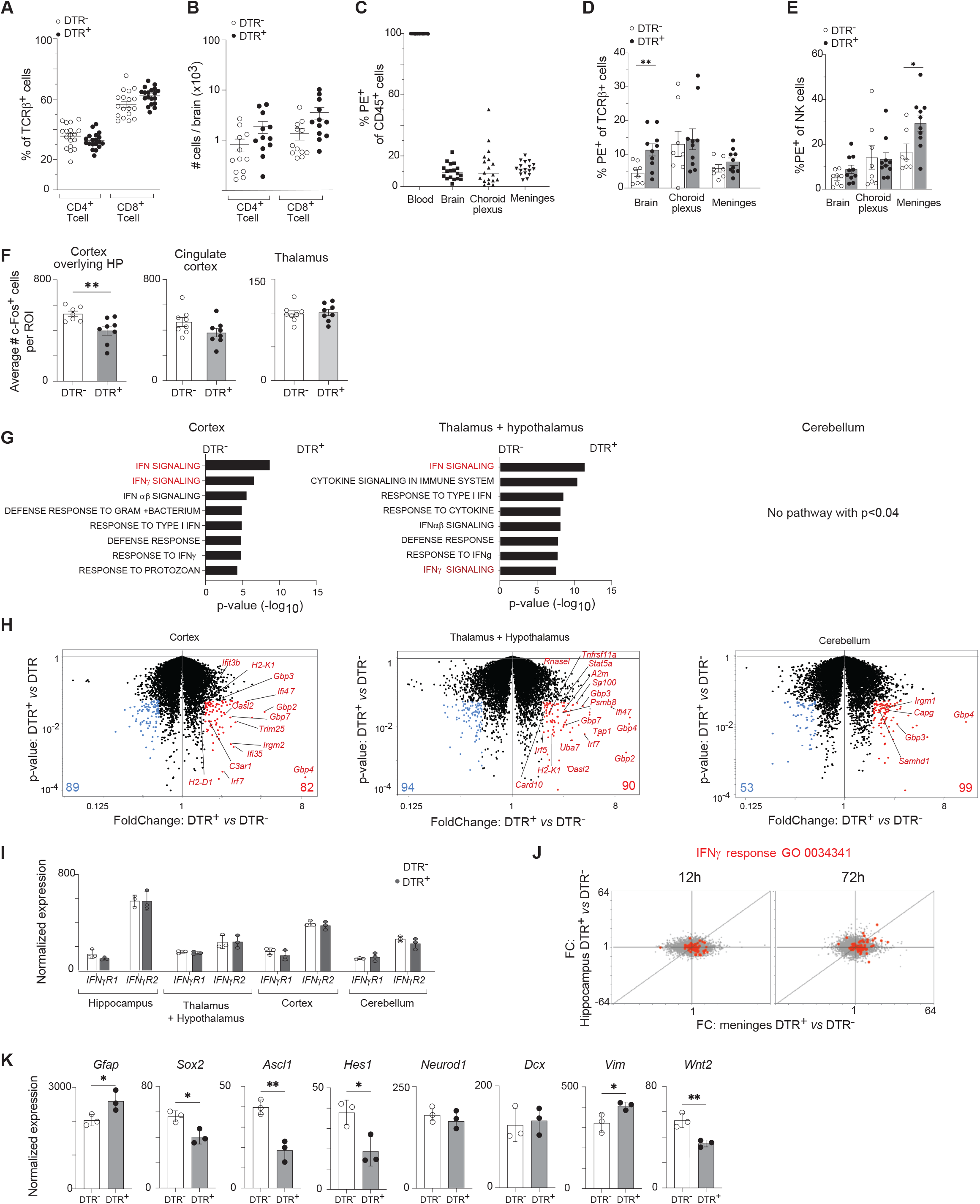
Additional characterization of the brain parenchyma response. DTR+ and DTR- mice were depleted of Treg cells as illustrated in Fig. 3A and were analyzed 5 days after the initial DT injection for changes in the brain parenchyma. (A-B) Flow cytometric analyses of CD4+ and CD8+ T cells in the brain parenchyma. (A) Frequency. n=12 (B) Number. n=12 (C-E) Flow cytometry analysis of Intravascular labelling of immunocytes with anti-CD45-PE mAb. (C) Total PE+ immunocytes in the indicated tissues. n=18 (D) Frequency of PE+ T cells. n≥8 (E) Frequency of PE+ NK cells. n≥8 (F) Histological quantification of cFos+ cells in the non-hippocampal brain regions mentioned in Fig. 4C. (G) Pathway-enrichment analyses (GO dataset and GSEA/MSigDB dataset) on the transcripts differentially expressed (FC>1.5, p<0.05) in H for the various brain regions in the presence vs absence of Treg cells. (H) Volcano plots annotated genes related to pathways highlighted in G (IFN signaling and IFNγ-response transcripts). Triplicate samples. (I) IFNγ receptor expression in the selected regions illustrated in Fig. 4G. RNA-seq data from Fig. 4G. (J) Fold-change/fold-change plot of whole-tissue hippocampal and meningeal transcriptomes in the absence vs presence of Treg cells 12h (left) and 72h (right) after the first DT injection. The IFNγ-response gene signature is highlighted in red. Duplicate samples. (K) Normalized hippocampal expression of key neurogenesis genes. RNA-seq data from Fig. 4H. PE, phycoerythrin; ROI, region of interest; HP, hippocampus, FC, fold-change. Mean ± SEM, except for (I) and (K) which show mean ± SD. P-values as per Fig. 3.

**Figure S7.**
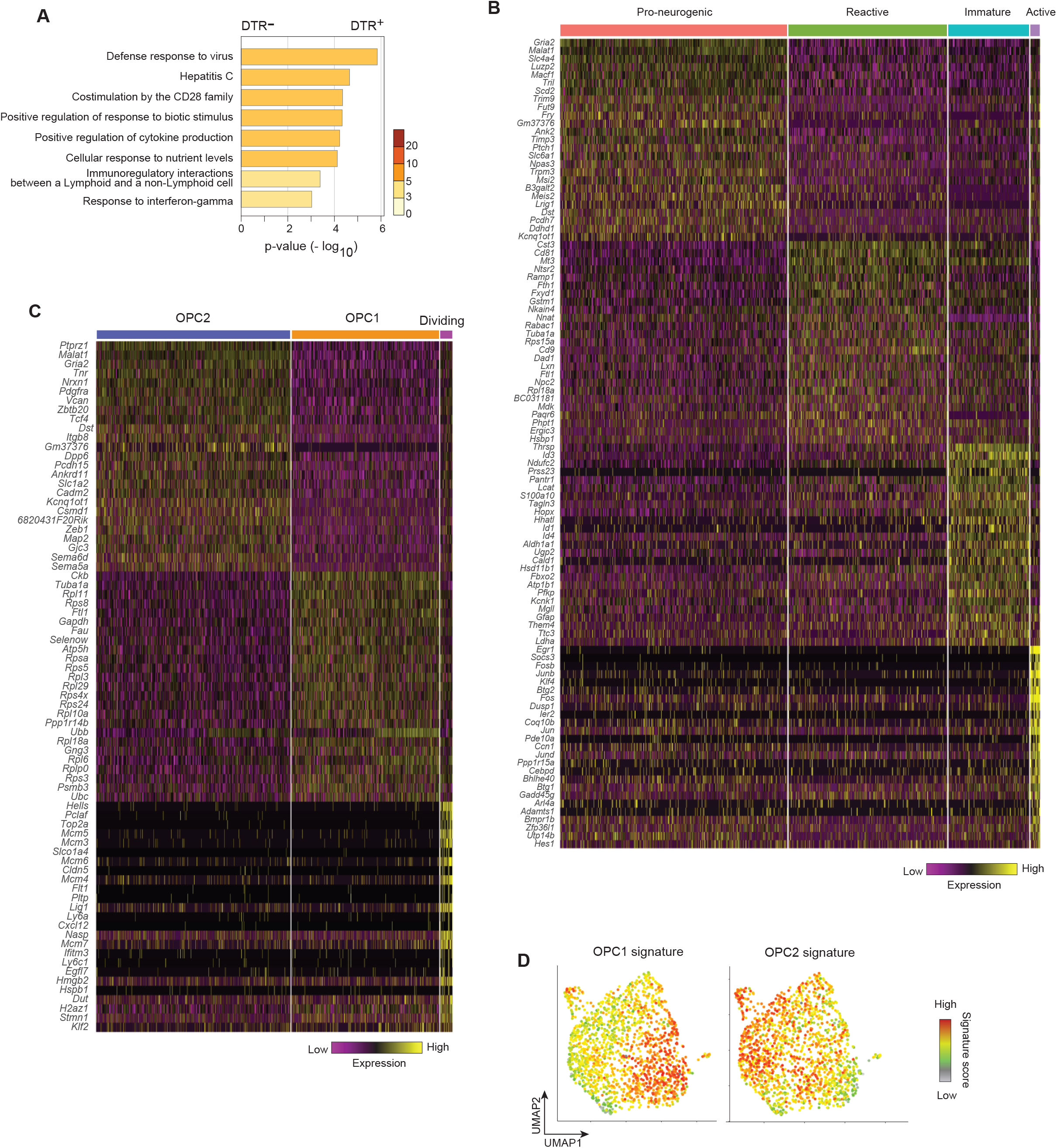
Additional characterization of the hippocampal glia populations. (A) Pathway enrichment analysis of the genes differentially expressed by (FC>1.5 p<0.05) microglia. Taken from Fig. 5B. (B) Heatmap of the 25 genes most differentially expressed by the astrocyte (light green) cluster of Fig. 5D. (C) Same as panel B, except for the OPC cluster. (D) Signature scores for the OPC1 (left) and OPC2 (right) subtypes (*74*) overlaid on the OPC UMAP of Fig. 5H. Abbreviations as per Fig. 5

**Figure S8.**
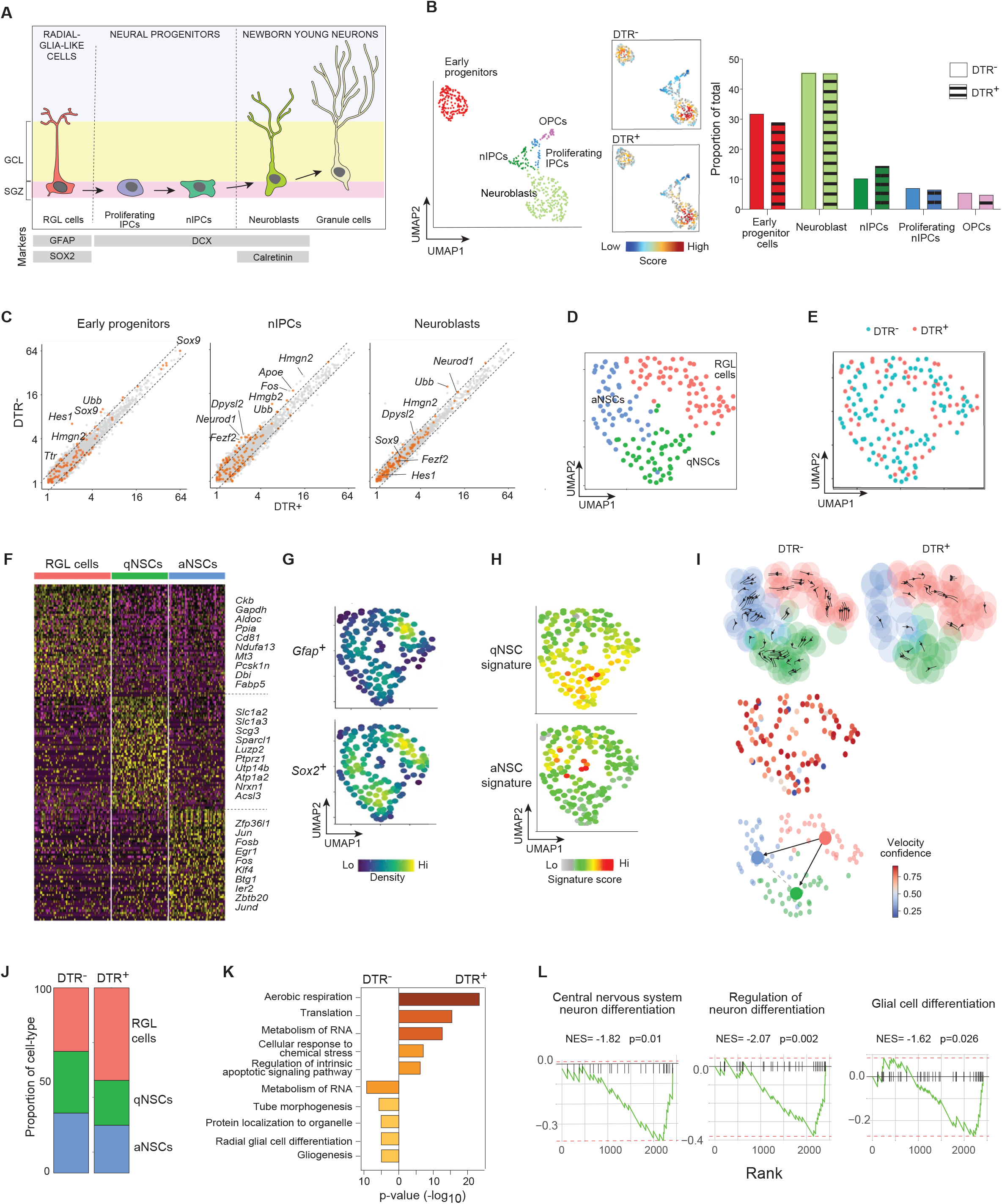
Regulation of hippocampal neurogenesis by Treg cells. (A) Simplified representation of neurogenesis in the DG of the hippocampus. The canonical markers that we have used to analyze each stage are annotated at the bottom. (B) Left panel: UMAP of reclustered neuronal populations from Fig. 5D; center, local densities of the various neuronal clusters in brains of mice with (top) or without (bottom) Treg cells; right: quantification of the various neuronal clusters. (C) Transcriptomic comparisons of the various neuronal clusters from mice with and without Treg cells. Relevant neurogenesis-related transcripts expressed differentially (FC>1.3) are annotated. (D) UMAP representation of the red clusters of early progenitor cells extracted from Fig. S8B. (E) UMAP representation of Fig. S8D comparing the distribution of cells from the DTR- (aquamarine) and DTR+ (coral) early progenitor cluster. (F) Heatmap of the 50 transcripts most differentially expressed by the three early progenitor subtypes, with the 10 most different listed. Abbreviations as per Fig. S8D. (G) Expression density plots of the two canonical RGL-cell markers overlain on the UMAP space of panel D. *Gfap* (top) and *Sox2* (bottom). (H) Activated and quiescent neuronal stem cell signature (*84*) scores overlain on the UMAP space of panel D. (I) Velocity trajectory analysis. Top row, summarized velocity vector field; middle row, velocity vector field coherence (confidence); bottom row, PAGA of the velocity trajectories across clusters. (J) Sub-type proportions within the early progenitor cluster. (K) Pathway-enrichment analysis via Metascape (*131*) of the RGL-cell cluster from panel D. (L) Relevant Gene-Set-Enrichment Analysis (GSEA) plots comparing differentially expressed RGL-cell transcripts in the presence vs absence of Treg cells. RGL, radial-glial-like; IPC, intermediate progenitor cells; nIPC, neuronal intermediate progenitor cells; qNSC, quiescent neuronal stem cell; aNSC, activated neuronal stem cell. Mean ± SEM, p-values as per Fig.3.

**Figure S9.**
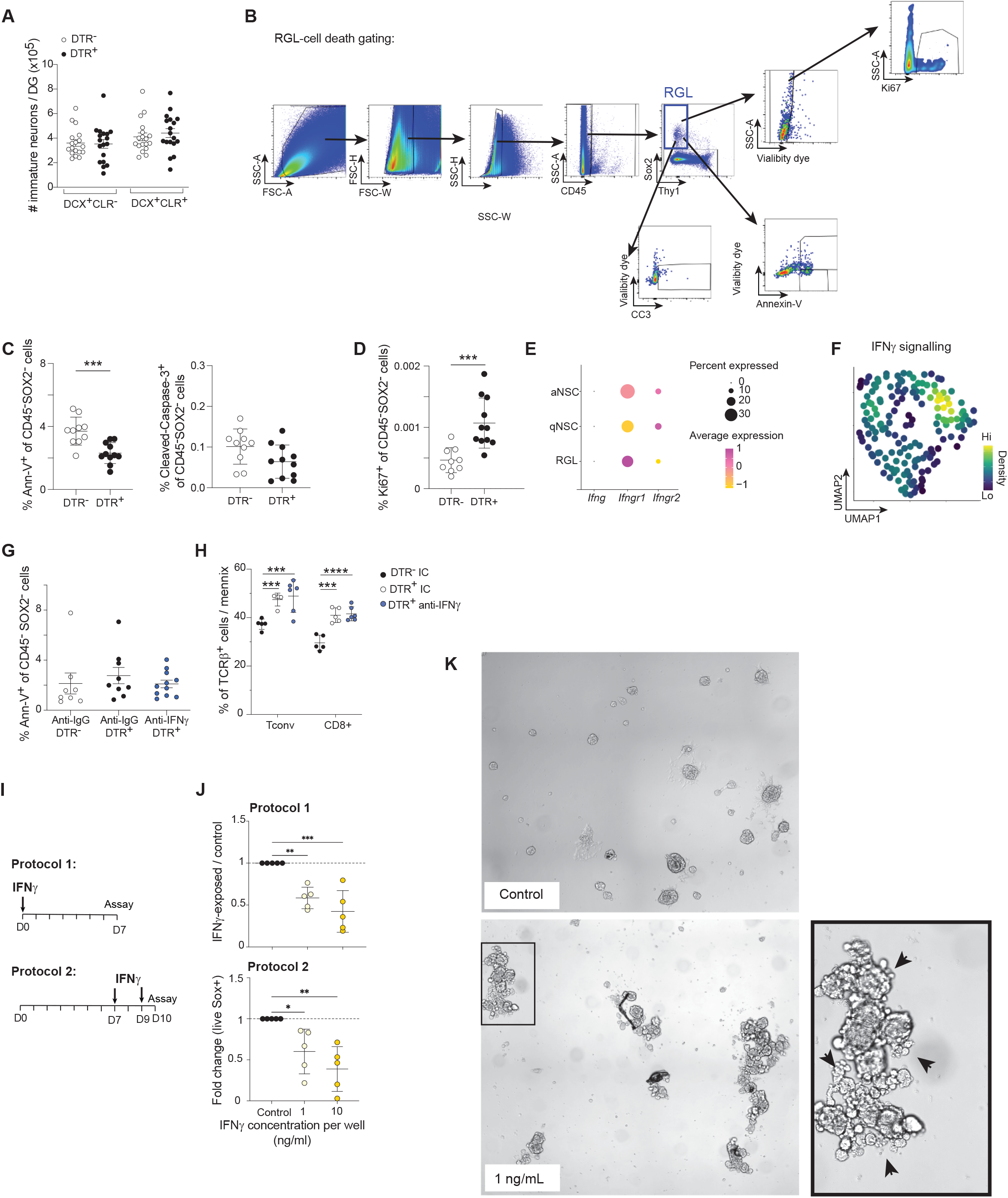
Additional characterization of the hippocampal neuronal clusters. (A) Histological quantification of two immature-neuron (DCX+) stages. n=18 (B) Gating strategy for the flow-cytometric identification of neural apoptosis (Annexin-V, CC3) and proliferation (ki67) in the hippocampus (C-D) Flow cytometric analysis of dying or cycling hippocampal CD45-SOX2-cells in the presence or absence of Treg cells. (C) Percentage of Annexin-V+ cells (left) and cleaved-caspase-3+ cells (right). n≥10 (D) Percentage of Ki67+ cells. n≥10 (E) Dot plot of *Ifnγ*, *Ifngr1*, and *Ifngr2* expression for each early progenitor cluster from Fig. S8D. (F) Expression density plot of “IFN gamma signaling” pathway from the GO database overlain on the UMAP space of Fig.S8D. (G) Quantification of Annexin-V+ cells in non-RGL cells (CD45-SOX2-) in DTR+ or DTR-mice with or without neutralization of IFNγ. n≥8 (H) Quantification of T cell subsets in the meninges in DTR+ or DTR- mice with or without neutralization of IFNγ. N=5 (I-K) Neurosphere cultures treated with rIFNγ at different stages of their generation. Neurospheres were generated by combining post-natal day 3 (P3) to post-natal day 10 (P10) hippocampi from male pups as specified in Methods. (I) Scheme for rIFNγ administration. (J) Total live SOX2+ cells after rIFNγ treatment compared to controls according to scheme on panel H. (K) Representative images of control (top) and rIFNγ-treated (bottom) neurosphere cultures from protocol 1. Arrows in the enlarged panel indicate apoptotic blebs on rIFNγ- treated neurospheres. CC3, cleaved-caspase 3. Mean ± SEM. p-values as per Fig. 1.

**Figure S10.**
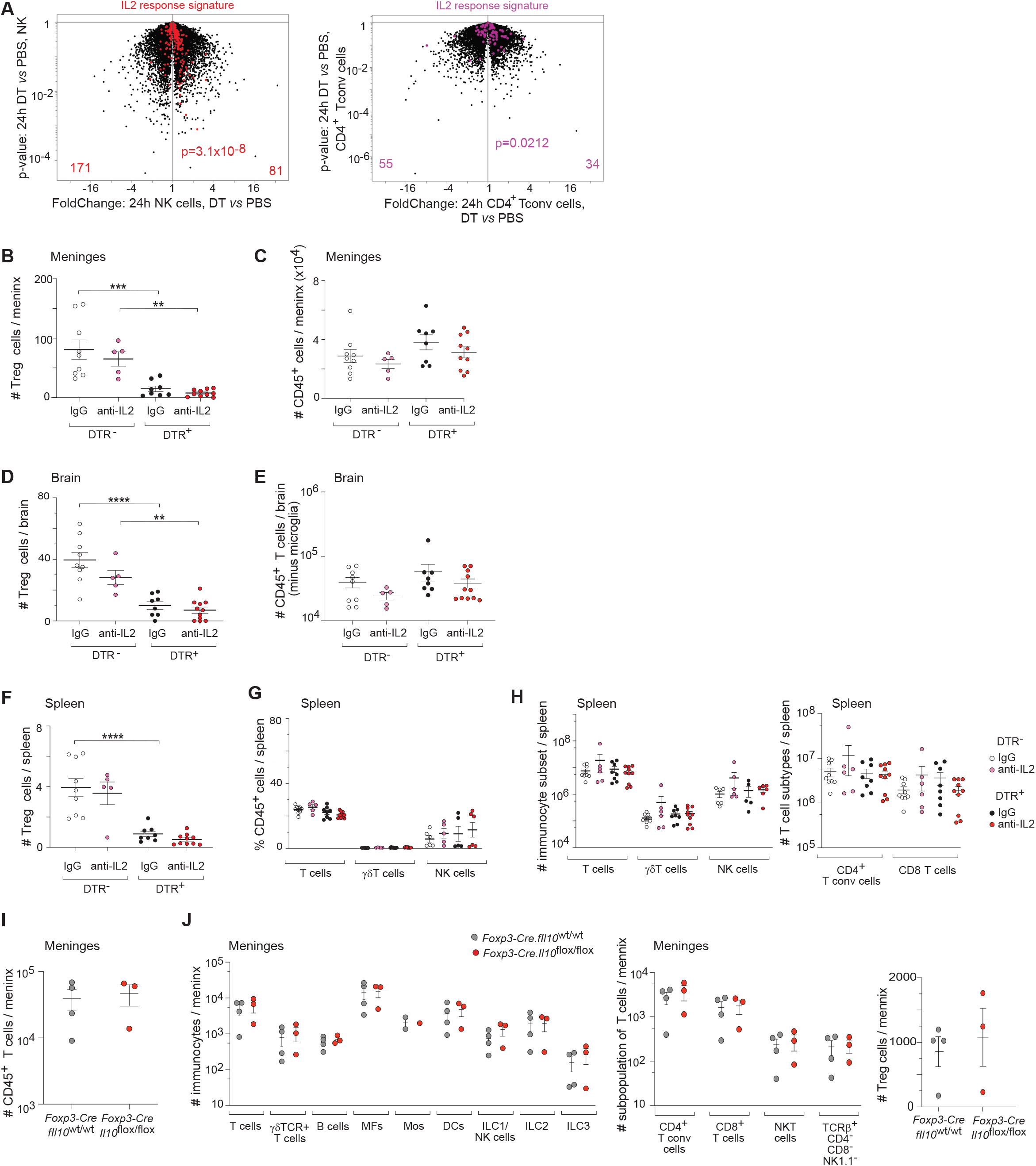
Additional characterization of the meningeal regulation of IL-2 levels by Treg cells. (A) “IL-2 response signature” (*135*) is highlighted on a volcano plot comparing transcripts expressed in sorted meningeal NK cells (left) and Tconvs (right) 24h after the first DT injection. Triplicate samples. (B-H) DTR+ and DTR- male littermates (6-7wks old) ip-injected with DT and anti-IL2 or isotype control mAb, as illustrated in Fig. 7B. n ≥ 5 B) Quantification of Treg cells in the meninges. (C) Quantification of total meningeal immunocytes. (D) Quantification of Treg cells in the brain parenchyma. (E) Quantification of total brain immunocytes. (F) Quantification of Treg cells in the spleen. (G) Percentage of select splenic immunocyte populations. (H) Numbers of splenic select immunocyte populations (left) and T cell subsets (right). (I-J) Flow-cytometric analysis of the dura mater from male littermates (6-10wks old) in which Treg cells express Il-10 (*Foxp3.Cre.Il10 wt/wt*) or lack Il-10 expression (*Foxp3.Cre.Il10 flox/flox*). n≥3 (I) Total meningeal immunocyte counts. (J) Quantification of select meningeal immunocyte populations (left), T cell subsets (center) and Treg cells (right). Mos, monocytes. Mean ± SEM. p-values as per Fig. 1. For simplicity, only relevant statistical differences are shown.

**Figure S11.**
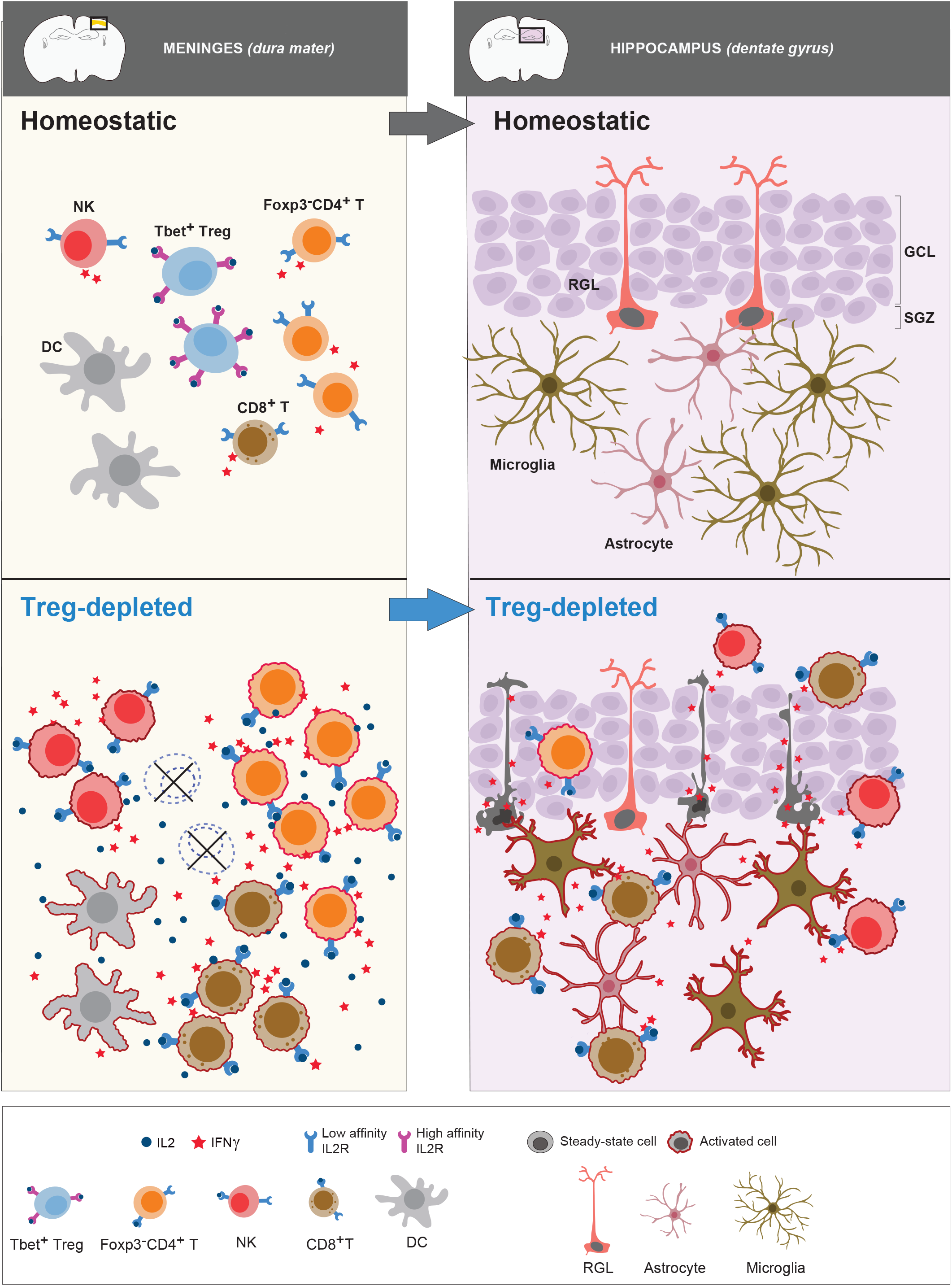
Schematic representation of the mechanism by which meningeal Tregs safeguard brain homeostasis in health individuals. Meningeal Tregs shield RGL cells by an indirect mechanism: their high-affinity IL-2 receptors sequester local IL-2, thereby starving local T and NK effector cells of a critical growth and differentiation factor, stifling their IFNγ production, and preventing their entry into the brain parenchyma, consequently, glial cells become activated, NSCs undergo IFNγ-dependent death, and short- and long-term spatial memory is compromised.

## SUPPLEMENTARY TABLE LEGENDS

**Table S1:** Repeated TCR sequences from Fig 2H.

**Table S2:** Summary and quality-control metrics of scRNA-seq datasets

