## Supplementary material for "The meninges host a unique compartment of regulatory T cells that bulwarks adult hippocampal neurogenesis": Table S2

**Table S2: Summary and quality-control metrics of scRNA-seq datasets**

**scRNA-seq/scTCR-seq**

| Dataset | Genotype + Treatment | Sorting strategy | Hashtaged | # mice | Total cells passing QC | Minimum genes / cell threshold | Minimum UMI / cell threshold | Maximum UMI /cell threshold | Maximum % mito / cell | Median genes / cell | Median UMI / cell | Median % mito / cell |
|---|---|---|---|---|---|---|---|---|---|---|---|---|
| Meningeal Tregs_Batch 1 | Foxp3Gfp | CD45+TCRβ+CD4+ | Yes (Hash 7) | 15 | 1154 | 500 | 500 | 4500 | < 6% | 1334 | 2971 | 2.03 |
| Meningeal Tregs_Batch 2 | B6.WT | CD45+TCRβ+CD4+ | Yes (Hash 10) | 25 | 2936 | 500 | 500 | 4500 | < 6% | 1094 | 2117 | 2.37 |

**scRNA-seq**

| Dataset | Genotype / Treatment | Sorting strategy | Hashtaged | # mice | Total cells passing QC | Minimum genes/cell threshold | Minimum UMI / cell threshold | Maximum UMI /cell threshold | Maximum % mito / cell | Median genes / cell | Median UMI / cell | Median % mito / cell |
|---|---|---|---|---|---|---|---|---|---|---|---|---|
| Hippocampus d5 post DT | DTR- / DT | CD45- Thy1- | No (SI-TT-B3) | 2 | 5509 | 500 | 500 | 1500 | 6 | 2286 | 6104 | 2.2 |
| Hippocampus d5 post DT | DTR+ / DT | CD45- Thy1- | No (SI-TT-A3) | 2 | 4725 | 500 | 500 | 1500 | 6 | 2061 | 4652 | 2.07 |
